## Supporting Information for "Evolutionarily related host and microbial pathways regulate fat desaturation"

### Materials and methods

#### General synthetic procedures.

Unless stated otherwise, all reactions were carried out under argon (Ar) atmosphere in flame-dried glassware. All commercially available reagents were used as purchased unless otherwise stated. All solvents were dried over activated 3Å molecular sieves for a minimum of 24 hours unless used in reactions containing aqueous reagents. Solutions and solvents sensitive to moisture and oxygen were transferred via standard syringe and cannula techniques. Reactions were cooled with ice-water or dry ice-acetone baths or heated with mineral oil baths depending on reaction temperature. Titanium (IV) isopropoxide was distilled under vacuum and stored under argon. Thin-layer chromatography (TLC) was performed with J.T. Baker Silica Gell IB2-F plastic-backed plates with analysis via UV and p-anisaldehyde stain. Flash chromatography was performed using Teledyne ISCO CombiFlash Rf and Rf+ systems with Teledyne ISCO RediSep Rf and Rf Gold silica columns. All deuterated solvents were purchased from Cambridge Isotopes. Nuclear Magnetic Resonance (NMR) spectra were recorded on a Varian INOVA 600 (600 MHz) or Bruker AV 500 (500 MHz) in the Cornell University NMR Facility. $^{1}H$ NMR chemical shifts are reported in ppm (δ) relative to the residual solvent peaks (7.26 ppm for $CDCl_3$ and 3.31 ppm for $CD_3OD$) and $^{13}C$ NMR shifts relative to their respective residual solvent peaks (77.16 for $CDCl_3$ and 49.00 for $CD_3OD$). NMR-spectroscopic data are reported as follows: chemical shift, multiplicity (s = singlet, d = doublet, t = triplet, q = quartet, m = multiplet, br = broad), coupling constants (Hz), and integration and often tabulated including 2D NMR data. All NMR data processing was done using MNOVA 12.0.1 (https://mestrelab.com/).

**Chemical syntheses**

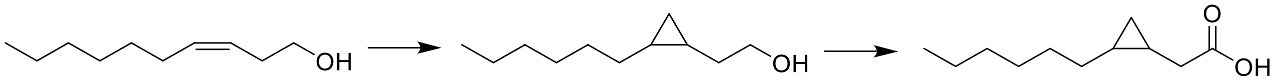

(**1**) (**2**) becyp#1

***cis*-3,4-methylenedecanoic acid, becyp#1.** From an 8:1 cis:trans solution of **1** (40 mg, 0.256 mmol, 1.00 equiv.) in 1.5 mL DCM was added $Zn(Et)_2$ (1M in hexanes, 1.28 mL, 1.28 mmol, 5.00 equiv.). The solution was cooled to 0 ˚C followed by the addition of $CH_2I_2$ (102 µL, 1.28 mmol, 5.00 equiv.). The resulting suspension was allowed to reach room temp with stirring and remained at room temp. overnight. The yellow mixture was directly purified via flash column chromatography on silica gel using a gradient of 5-100% EtOAc in hexanes, affording cyclopropyl alcohol intermediate (**2**, 40 mg) with some uncharacterized impurities. To a solution of the cyclopropyl alcohol intermediate (**2**, 10 mg) in 1 mL of acetone at 0 ˚C was added 6 drops of freshly prepared Jones reagent (enough to maintain an orange color). The solution was stirred at 0 ˚C for 1 hr and then directly purified via flash column chromatography on silica gel using a gradient of 5-100% EtOAc in hexanes, affording becyp#1 (8 mg, 69% over two steps) as a colorless oil, determined as a mixture of 8:1 *cis : trans* isomers.

**$^1$H NMR (600 MHz, chloroform-*d*):** δ 2.42 (dd, *J* = 16.6, 7.0 Hz, 1H), 2.30 (dd, *J* = 16.6, 7.6 Hz, 1H), 1.44 – 1.20 (m, 10H), 1.15 (m, 1H), 1.09 (m, 1H), 0.88 (t, *J* = 6.8 Hz, 3H), 0.74 (td, *J* = 8.4, 4.6 Hz, 1H), -0.11 (q, *J* = 5.2 Hz, 1H). These chemical shifts were nearly identical to those previously reported[1].

**$^{13}$C NMR (126 MHz, chloroform-*d*):** δ 180.1, 33.8, 32.2, 30.0, 29.4, 28.9, 22.8, 15.6, 14.2, 11.3, 11.

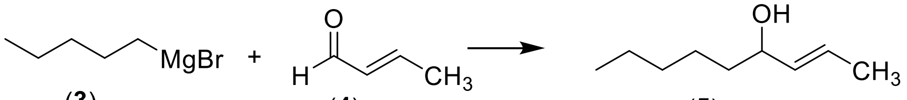

(**3**) (**4**) (**5**)

***(E)*-4-Hydroxy-non-2-ene (5).** *n*-Pentylmagnesium bromide (**3**, 10 mL, 20 mmol in THF) was added to a diethyl ether (20mL) at 0 °C under an argon atmosphere. Crotonaldehyde (**4**, 1.66 mL, 20 mmol) was added to the stirring solution over 10 minutes via an addition funnel and stirred for one hour. The reaction was quenched with saturated aqueous NH4Cl (20 mL) and extracted with EtOAc (3×20 mL). The combined organics were dried over $MgSO_4$, filtered, and concentrated under reduced pressure. The resulting liquid was purified by flash chromatography on silica gel. Elution with a gradient of 0-20% EtOAc/Hexanes gave the resulting alcohol (**5**) (2.61 g, 92%) as a colorless liquid[2].

**$^1$H NMR (CDCl3, 500 MHz):** δ (ppm) 5.65 (dqd 15.3, 6.5, 0.8 Hz, 1H), 5.48 (ddq 15.3, 7.7, 1.5 Hz, 1H), 4.03 (q, 6.6 Hz, 1H), 1.70 (dd 6.5, 1.3 Hz, 3H), 1.25-1.58 (m, 9H), 0.89 (t, 6.8 Hz, 3H).
**$^{13}$C NMR (CDCl3, 125 MHz):** δ (ppm) 134.5, 126.9, 73.4, 37.4, 31.9, 25.3, 22.8, 17.8, 14.2.
**HRMS (ESI) *m/z*:** Calculated: $(M+H)^+$ 143.1430. Actual: 143.1428 Δ ppm: -1.64

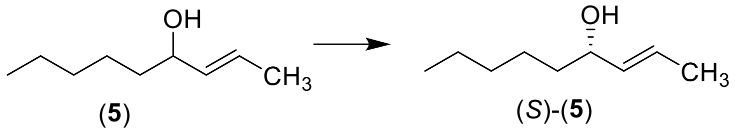

***(S,E)*-4-Hydroxy-non-2-ene (*(S)*-5).** *(-)*-Diisopropyl D-tartrate (183 µL, 0.88 mmol) was added to a suspension of powdered 4 Å molecular sieves (0.4 g) in DCM (21 mL) at ambient temperature under an argon atmosphere. The solution was cooled to -20 °C and titanium (IV) isopropoxide (212 µL, 0.7 mmol) was added. The solution was stirred for one hour and *tert*-butylhydroperoxide (382 µL, 2.1 mmol in decane) added. The solution was further stirred for 30 minutes and cooled to -40 °C, upon which racemic *(E)*-4-hydroxy-non-2-ene (**5**) (3.5 mL, 3.5 mmol in DCM) was added dropwise. After 20 hours the reaction was quenched with *(-)*-diisopropyl D-tartrate (366 µL, 1.75 mmol) in water (7 mL). The layers were separated, and the aqueous solution extracted with $Et_2O$ (3×20 mL). The combined organics were washed with saturated aqueous $NaHCO_3$, dried over $MgSO_4$, filtered, and concentrated under reduced pressure. The resulting liquid was purified by flash chromatography on silica gel as above, yielding chiral alcohol *(S)*-**5** (236 mg, 94%) as a colorless liquid[3]. Enantiomeric excess was determined to be up to 88% by Mosher derivatization[4]. All NMR spectra and mass spectrometric data are identical to racemic alcohol (**5**).

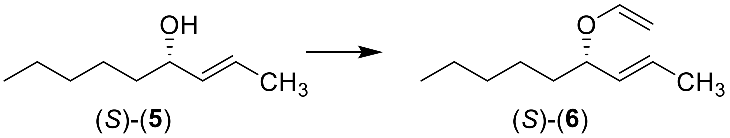

***(S,E)*-4-(Vinyloxy)-non-2-ene (*(S)*-6).** *(S,E)*-4-Hydroxy-non-2-ene (*(S)*-**5**, 307 mg, 2.16 mmol) was added to ethyl vinyl ether (5.75 mL, 60.5 mmol) at ambient temperature under an argon atmosphere. Mercury (II) acetate (688 mg, 2.16 mmol) was added to the solution[5]. After two hours AcOH (302 µL) was added with stirring. After 30 minutes the reaction was diluted with hexanes (45 mL) and washed with 5% aqueous KOH (4.5 mL). The organics were dried over $MgSO_4$, filtered, and concentrated under reduced pressure. The resulting liquid was purified by flash chromatography on alumina. Elution with a gradient of 0-10% EtOAc/Hexanes yielded vinyl ether (*(S)*-**6**, 178 mg, 76% BRSM) as a colorless liquid.

**$^1$H NMR ($CDCl_3$, 500 MHz)** δ (ppm) 6.31 (dd, 14.1, 6.6 Hz, 1H), 5.65 (dqd 15.3, 6.6, 0.8 Hz, 1H), 5.38 (ddq, 15.3, 7.6, 1.6 Hz, 1H), 4.28 (dd, 14.1, 1.4 Hz, 1H), 4.05 (q, 7 Hz, 1H), 3.96 (dd, 6.6, 1.4 Hz, 1H), 1.71 (dd, 6.5, 1.4 Hz, 3H), 1.61-1.70 (m, 1H), 1.46-1.54 (m, 1H), 1.23-1.40 (m, 6H), 0.88 (t, 6.8 Hz, 3H).
**$^{13}$C NMR ($CDCl_3$, 125 MHz)** δ (ppm) 151.0, 131.4, 128.7, 88.4, 81.2, 35.3, 31.8, 25.0, 22.7, 17.9, 14.2.
**HRMS (ESI) *m/z*:** Calculated: $(M+H)^+$ 169.1587 $(M+Na)^+$ 191.1406. Actual: 169.1584 Δ ppm: -1.99

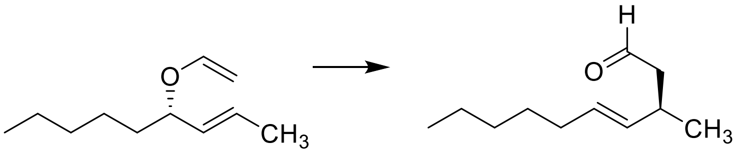

(*S*)-(**6**) (*R*)-(**7**)

***(R,E)*-3-Methyl-dec-4-enal (*(R)*-7).** Vinyl ether (*(S)*-**6**, 107 mg, 0.64 mmol) was added to toluene (5 mL) and stirred with condenser at reflux under an argon atmosphere. After 23 hours the reaction was concentrated to yield aldehyde (*(R)*-**7**, 100 mg, 93%) as a colorless liquid which was used without purification[5–7].
**$^1$H NMR (CDCl3, 500 MHz)** δ (ppm) 9.72 (t, 2.4 Hz, 1H), 5.44 (dtd, 15.4, 6.6, 1.0 Hz, 1H), 5.34 (ddt, 15.4, 7.0, 1.3 Hz, 1H), 2.72 (m, 6.9 Hz, 1H), 2.40 (ddd, 16.0, 7.3, 2.4 Hz, 1H), 2.33 (ddd, 16.0, 6.7, 2.4 Hz, 1H), 1.97 (q, 7.0 Hz, 2H), 1.21-1.37 (m, 6H), 1.06 (d, 6.8 Hz, 3H), 0.88 (t, 7.0 Hz, 3H).
**$^{13}$C NMR (CDCl3, 125 MHz)** δ (ppm) 203.0, 133.9, 130.2, 50.7, 32.6, 31.8, 31.5, 29.3, 22.7, 20.9, 14.2.
**HRMS (ESI) *m/z*:** Calculated: $(M+Na)^+$ 191.1406. Actual: 191.1411 Δ ppm: 2.49

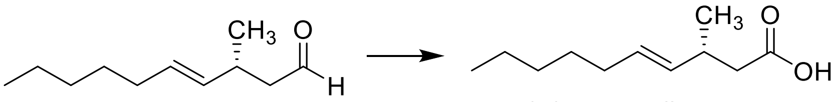

(*R*)-(**7**) (*R*)-bemeth#1

***(R,E)*-3-Methyl-dec-4-enoic acid, *(R)*-bemeth#1.** Aldehyde (*(R)*-**13**, 53 mg, 0.32 mmol) was dissolved in DMSO (1.1 mL) and stirred under ambient atmosphere. Sodium chlorite (40 mg, 0.35 mmol) was dissolved in minimal water and the pH adjusted to ~4.5 with $NaH_2PO_4$. The aqueous solution was added to the aldehyde and the reaction stirred in an open atmosphere. After 30 minutes, more sodium chlorite was added to the reaction (20 mg, 0.18 mmol) dissolved in water and buffered as above. After 45 minutes the reaction was diluted with water (2 mL) and extracted with EtOAc (3×2 mL). The combined organics were dried over $MgSO_4$, filtered, and concentrated under reduced pressure. The resulting oil was purified by flash chromatography on silica gel. Elution with a gradient of 0-20% EtOAc(0.1% AcOH)/hexanes yielded *(R)*-bemeth#1 (**4**, 44 mg, 76%) as a colorless oil. Enantiomeric excess was determined to be 65% by 2,2,2- trifluoro-1-phenethylamine derivatization and subsequent analysis by UHPLC-MS.

**$^{1}$H NMR (CDCl3, 500 MHz)** δ (ppm) 5.45 (dtd, 15.4, 6.7, 0.8 Hz, 1H), 5.33, ddt, 15.3, 7.2, 1.3, 1H), 2.63 (m, 7.0 Hz, 1H), 2.35 (dd, 14.9, 7.3 Hz, 1H), 2.28 (dd, 14.9, 7.3 Hz, 1H), 1.96 (q, 7.0, 2H), 1.21-1.36 (m, 6H), 1.05 (d, 6.7 Hz, 3H), 0.88 (t, 6.9 Hz, 3H).

**$^{13}$C NMR (CDCl3, 125 MHz)** δ (ppm) 178.8, 133.7, 1.0.1, 41.8, 33.5, 32.6, 34.4, 29.3, 22.7, 20.5, 14.2.

**HRMS (ESI) *m/z*:** Calculated: $(M-H)^-$ 183.1391. Actual: 183.1383. Δ ppm: -4.16.

***(S,E)*-3-Methyl-dec-4-enoic acid, *(S)*-bemeth#1.** Following Sharpless resolution using *(+)*-Diisopropyl L-tartrate, parallel reactions yielded *(S)*-bemeth#1 with identical physical and spectroscopic properties.

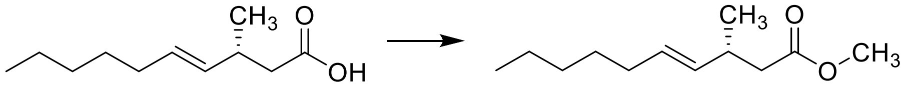

(*R*)-bemeth#1 (*R*)-(**8**)

**Methyl-*(R, E)*-3-methyl-dec-4-enoate (*(R)*-8).** Trimethylsilyldiazomethane (1.2 mL, 0.72 mmol) was added dropwise to a solution of *(R)*-bemeth#1 in DCM (3 mL) and MeOH (3 mL). After stirring for 15 minutes, the reaction was concentrated under reduced pressure. The resulting oil was purified by flash chromatography on silica gel. Elution with a gradient of 0- 10% EtOAc/hexanes yielded methyl ester (*(R)*-**8**, 42 mg, 89%) as a colorless oil.

**$^{1}$H NMR (CDCl3, 500 MHz)** δ (ppm) 5.42 (dtd, 15.3, 6.7, 0.9 Hz, 1H), 5.31 (ddt, 15.3, 7.3, 1.2 Hz, 1H), 3.63 (s, 3H), 2.61 (m, 7.0 Hz, 1H), 2.30 (dd, 14.7, 7.3 Hz, 1H), 2.24 (dd, 14.7, 7.3 Hz, 1H), 1.95 (q, 6.9 Hz, 2H), 1.21-1.35 (m, 6H), 1.02 (d, 6.9 Hz, 3H), 0.87 (t, 6.9 Hz, 3H).

**$^{13}$C NMR (CDCl3, 125 MHz)** δ (ppm) 173.3, 134.0, 129.8, 51.5, 42.0, 33.8, 32.6, 31.4, 29.3, 22.7, 20.6, 14.

**HRMS (ESI) *m/z*:** Calculated: $(M+H)^+$ 199.1693. Actual: 199.1690. Δ ppm: -1.49.

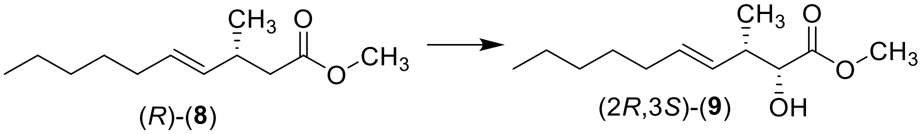

(*R*)-(**8**) (2*R*,3*S*)-(**9**)

**Methyl-*(2R,3S,E)*-2-hydroxy-3-methyl-dec-4-enoate (*(2R,3S)*-9).** *n*-Butyl lithium (72 μL, 0.18 mmol) was added dropwise to a stirring solution of *N,N*-diisopropylamine (21 μL, 0.15 mmol) in THF (4 mL) at -15°C and stirred 10 minutes under an argon atmosphere. The solution was cooled to -78°C and methyl ester (*(R)*-**8**) was added and the reaction stirred at -15°C for 15 minutes. The reaction was cooled to -78°C and *(+)*-(8, 8-dichlorocamphorylsulfonyl) oxaziridine (90 mg, 0.3 mmol) was added in THF (2 mL) and stirred to -15°C[8]. After stirring for one hour, the reaction was quenched with aqueous saturated $NaHCO_3$ (3 mL) and the layers separated. The aqueous layer was extracted with DCM (3×5mL) and the combined organics dried over $MgSO_4$, filtered, and concentrated under reduced pressure. The resulting oil was purified by flash chromatography on silica gel. Elution with a gradient of 0-20% EtOAc/hexanes yielded alpha-hydroxy ester (*(2R,3S)*-**9**, 9 mg, 69% BRSM).
**$^1$H NMR (CDCl3, 500 MHz)** δ (ppm) 5.53 (dtd, 15.3, 6.7, 0.9 Hz, 1H), 5.39 (ddt, 15.3, 7.9, 1.4 Hz, 1H), 4.12 (d, 4.2, 1H), 3.78 (s, 3H), 3.76 (q, 4.3 Hz, 1H), 2.52-2.67 (m, 1H), 2.00 (qd, 7.1, 1.2 Hz, 2H), 1.21-1.38 (m, 8H), 0.99 (d, 7.0, 3H), 0.88 (t, 6.9, 3H).
**$^{13}$C NMR (CDCl3, 125 MHz)** δ (ppm) 174.8, 132.4, 130.6, 74.6, 52.4, 41.3, 32.7, 31.5, 29.2, 22.7, 14.8, 14.2.
**HRMS (ESI) *m/z*:** Calculated: $(M+Na)^+$ 237.1461. Actual: 237.1475. Δ ppm: 5.87.

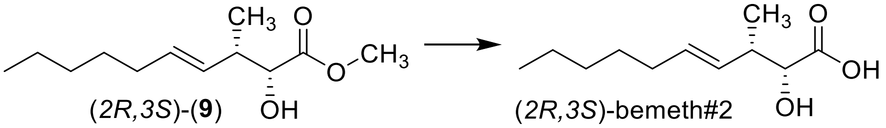

(*2R,3S*)-(**9**) (*2R,3S*)-bemeth#2

***(2R,3S,E)*-2-hydroxy-3-methyl-dec-4-enoic acid, *(2R,3S)*-bemeth#2.** Lithium hydroxide (40 mg, 2 mmol) was added to a stirring solution of ester (*(2R,3S)*-**9**, 11 mg, 0.05 mmol) in MeOH (0.4 mL), THF (0.4 mL), and water (0.2 mL). After one hour, the reaction was acidified with 1 M HCl and extracted with DCM (3×5 mL). The combined organics were dried over MgSO4, filtered, and concentrated under reduced pressure. The resulting oil was purified by flash chromatography on silica gel. Elution with a gradient of 0-100% DCM/MeOH(0.1% AcOH) yielded alpha-hydroxy acid *(2R,3S)*-bemeth#2 (8.6 mg, 86% BRSM, d.r. 67.8% as determined by Mosher analysis[9]).
**$^1$H NMR (CDCl3, 500 MHz)** δ (ppm) 5.58 (dt, 15.4, 6.8 Hz, 1H), 5.42 (dd, 15.4, 7.6 Hz, 1H), 4.22 (d, 3.6, 1H), 2.68 (m, 1H), 2.02 (q, 6.9 Hz, 2H), 1.22-1.40 (m, 6H), 1.05 (d, 7.0, 3H), 0.89 (t, 6.7, 3H).

**$^{13}$C NMR (CDCl3, 125 MHz)** δ (ppm) 177.4, 133.1, 130.1, 74.2, 40.7, 32.7, 31.5, 29.2, 22.7, 14.3, 14.2

**HRMS (ESI) *m/z*:** Calculated: $(M-H)^-$ 199.1340. Actual: 199.1339. Δ ppm: -0.28.

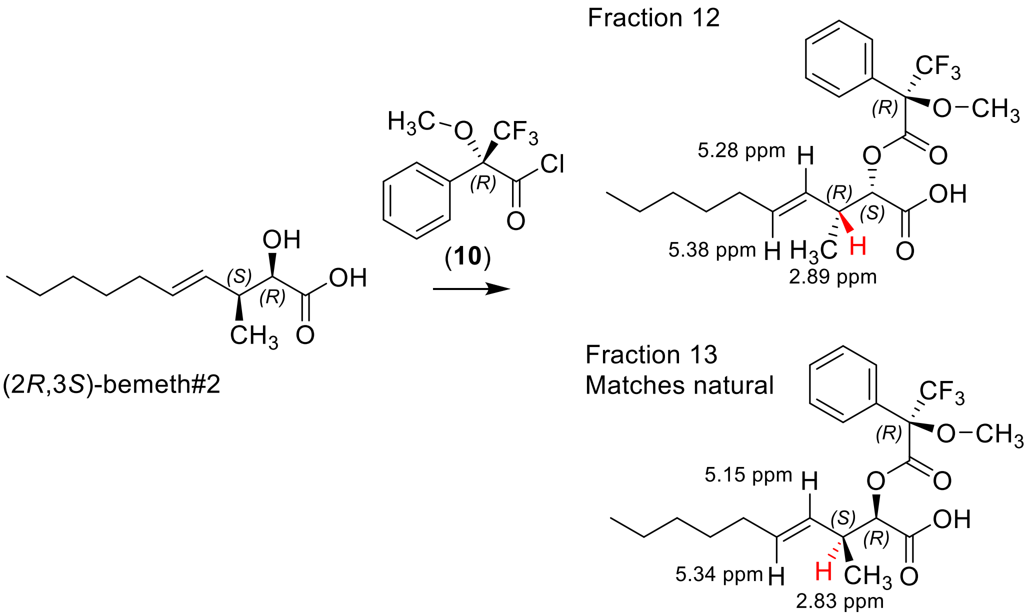

**Determination of bemeth#2 stereochemistry.** bemeth#2 was dissolved in DCM with DMAP and stirred under argon at ambient temperature. *(R)*-(+)-α-Methoxy-α-(trifluoromethyl)phenylacetyl chloride (**10**, 1.2 equivalents) was added and the reaction stirred for 30 minutes and quenched with MeOH. The reaction was concentrated under reduced pressure, taken up in MeOH, and analyzed by HPLC-HRMS. Integration of the EIC after Mosher derivatization of synthetic bemeth#2 yielded a diastereomeric enrichment of 68%.

### Supplementary Tables

**Supplementary Table 1. Metabolites enriched in *acdh-11(n5878)* mutants.** Subset of dereplicated metabolites that are i) at least 8-fold enriched in *acdh-11(n5878);P$_{fat-7}$::fat-7::GFP* relative to WT *P$_{fat-7}$::fat-7::GFP,* ii) mean intensity > 500,000 AU for *acdh-11*, iii) not detected in bacteria only, and iv) dependent on cyclopropane lipid biosynthesis in *E. coli*. These data were filtered using stringent criteria and hundreds of additional differential features were detected (not dereplicated) using mean intensity cutoff at 100,000 AU. Note: these metabolites were detected in *acdh-11(n5878)* animals reared on OP50, HB101, and BW25113.

| **ES(-) Obs. *m/z*** | **RT (min)** | **Molecular Formula** | **ES(-) Theor. *m/z*** | ***m/z* error (ppm)** | **SMID-DB #** | **Comments** |
|---|---|---|---|---|---|---|
| 242.10394 | 6.90 | $C_{11}H_{17}NO_5$ | 242.10340 | 2.226 | | |
| 171.06656 | 6.96 | $C_8H_{12}O_4$ | 171.06628 | 1.592 | | likely dicarboxylate, Na adduct in ES(-) |
| 457.14896 | 7.14 | $C_{17}H_{31}O_{12}P$ | 457.14804 | 2.013 | | endo |
| 574.20697 | 7.19 | $C_{25}H_{38}NO_{12}P$ | 574.20589 | 1.881 | | endo |
| 576.22237 | 7.32 | $C_{25}H_{40}NO_{12}P$ | 576.22154 | 1.449 | | endo |
| 396.17978 | 7.47 | $C_{16}H_{32}NO_8P$ | 396.17928 | 1.261 | | endo |
| 427.13823 | 7.62 | $C_{16}H_{28}O_{11}P$ | 427.13747 | 1.769 | | |
| 215.12915 | 7.75 | $C_{11}H_{20}O_4$ | 215.12888 | 1.225 | becyp#32 | |
| 229.10845 | 7.85 | $C_{11}H_{18}O_5$ | 229.10815 | 1.331 | | |
| 215.12920 | 7.94 | $C_{11}H_{20}O_4$ | 215.12888 | 1.483 | becyp#33 | |
| 185.08227 | 7.95 | $C_9H_{14}O_4$ | 185.08193 | 1.797 | | likely dicarboxylate, Na adduct in ES(-) |
| 227.09299 | 8.02 | $C_{11}H_{16}O_5$ | 227.09250 | 2.131 | | |
| 213.11365 | 8.12 | $C_{11}H_{18}O_4$ | 213.11323 | 1.979 | | *endo* |
| 576.22271 | 8.15 | $C_{25}H_{40}NO_{12}P$ | 576.22154 | 2.029 | | *endo* |
| 213.11378 | 8.16 | $C_{11}H_{18}O_4$ | 213.11323 | 2.591 | | |
| 215.12918 | 8.18 | $C_{11}H_{20}O_4$ | 215.12888 | 1.360 | becyp#3 | |
| 213.11380 | 8.18 | $C_{11}H_{18}O_4$ | 213.11323 | 2.662 | | |
| 439.13818 | 8.27 | $C_{17}H_{28}O_{11}P$ | 439.13747 | 1.600 | | *endo*, MOGL |
| 229.10852 | 8.30 | $C_{11}H_{18}O_5$ | 229.10815 | 1.630 | | |
| 286.16662 | 8.31 | $C_{14}H_{25}NO_5$ | 286.16600 | 2.179 | | |
| 441.15367 | 8.35 | $C_{17}H_{30}O_{11}P$ | 441.15312 | 1.239 | | *endo, MOGL* |
| 286.16672 | 8.38 | $C_{14}H_{25}NO_5$ | 286.16600 | 2.528 | | formate adduct |
| 276.13761 | 8.38 | $C_{13}H_{24}NO_3Cl$ | 276.13720 | 1.509 | | |
| 288.18226 | 8.41 | $C_{14}H_{27}NO_5$ | 288.18165 | 2.143 | | formate |

| | | | | | | |
|---|---|---|---|---|---|---|
| | | | | | | adduct |
| 213.11378 | 8.47 | $C_{11}H_{18}O_4$ | 213.11323 | 2.591 | | *endo* |
| 591.17201 | 8.49 | $C_{22}H_{32}N_4O_{13}P$ | 591.1708979 | 1.878 | | *endo* |
| 441.15376 | 8.54 | $C_{17}H_{30}O_{11}P$ | 441.15312 | 1.451 | | *endo* |
| 254.14035 | 8.61 | $C_{13}H_{21}NO_4$ | 254.13978 | 2.243 | | |
| 405.17729 | 8.62 | $C_{18}H_{29}O_{10}$ | 405.17662 | 1.652 | | formate adduct |
| 256.15597 | 8.63 | $C_{13}H_{23}NO_4$ | 256.15543 | 2.092 | | |
| 361.18767 | 8.70 | $C_{17}H_{30}O_8$ | 361.18680 | 2.444 | | |
| 199.09793 | 8.88 | $C_{10}H_{16}O_4$ | 199.09758 | 1.747 | | likely dicarboxylate, Na adduct in ES(-) |
| 716.30686 | 9.12 | $C_{33}H_{52}NO_{14}P$ (?) | 716.30527 | 2.226 | | endo |
| 270.17162 | 9.13 | $C_{14}H_{25}NO_5$ | 270.17108 | 1.999 | | |
| 758.35478 | 9.42 | $C_{36}H_{57}NO_{14}P$ (?) | 758.35222 | 3.383 | | endo |
| 756.33820 | 9.44 | $C_{36}H_{55}NO_{14}P$ | 756.33657 | 2.165 | | endo |
| 560.22734 | 9.58 | $C_{25}H_{40}NO_{11}P$ | 560.22662 | 1.285 | | endo |
| 199.13434 | 9.77 | $C_{11}H_{20}O_3$ | 199.13397 | 1.850 | becyp#2 | characterized by 2D-NMR |
| 213.11365 | 9.79 | $C_{11}H_{18}O_4$ | 213.11323 | 1.970 | becyp#4 | likely dicarboxylate, Na adduct in ES(-) |
| 560.19103 | 9.89 | $C_{24}H_{36}NO_{12}P$ | 560.19024 | 1.420 | | MOGL |
| 197.11851 | 9.90 | $C_{11}H_{18}O_3$ | 197.11832 | 0.988 | | endo |
| 558.17604 | 9.93 | $C_{24}H_{34}NO_{12}P$ | 558.17459 | 2.603 | | |
| 542.23839 | 9.96 | $C_{22}H_{42}NO_{12}P$ | 542.23719 | 1.864 | | GLEA |
| 199.13420 | 10.02 | $C_{11}H_{20}O_3$ | 199.13397 | 1.183 | becyp#22 | endo, minor |
| 639.27995 | 10.36 | $C_{28}H_{48}O_{14}P$ | 639.27871 | 1.928 | | endo |
| 411.14319 | 10.47 | $C_{16}H_{28}O_{10}P$ | 411.14255786 | 1.533 | | endo |
| 560.19090 | 10.57 | $C_{24}H_{36}NO_{12}P$ | 560.19024 | 1.179 | | endo, MOGL, anthranilate |
| 558.17534 | 10.66 | $C_{24}H_{34}NO_{12}P$ | 558.17459 | 1.345 | | endo |
| 584.19113 | 10.68 | $C_{26}H_{35}NO_{12}P$ | 584.19023 | 1.537 | | endo |
| 540.20149 | 10.74 | $C_{25}H_{36}NO_{10}P$ | 540.20041 | 2.009 | iglu#202 | MOGL |
| 411.14318 | 10.80 | $C_{16}H_{28}O_{10}P$ | 411.14256 | 1.512 | | endo |
| 377.18250 | 11.16 | $C_{17}H_{30}O_9$ | 377.18171 | 2.100 | | formate adduct |
| 544.19667 | 12.58 | $C_{24}H_{36}NO_{11}P$ (?) | 544.19532 | 2.468 | | |
| 335.17716 | 17.31 | $C_{21}H_{24}N_2O_2$ | 335.17650 | 1.950 | | |
| 183.13906 | 9.95 | $C_{11}H_{20}O_2$ | 183.13905 | 0.037 | becyp#1 | post-column ion pairing |

**Supplementary Table 2. FCMT-1-derived metabolites.** A comprehensive list of FCMT-1-derived metabolites that were detected in the conditioned media (*exo*-) or worm body *(endo*-) metabolome of N2 (WT) synchronized gravid adults in liquid culture, unless otherwise indicated.

| ES(-) Obs. *m/z* | RT (min) | Molecular Formula | ES(-) Theor. *m/z* | *m/z* error (ppm) | SMID-DB # | Comments |
|---|---|---|---|---|---|---|
| 231.12393 | 7.18 | $C_{11}H_{20}O_5$ | 213.12380 | 0.584 | bemeth#401 | enriched *endo*, H/D exchange $D_8/D_9/D_{10}$ |
| 215.12902 | 8.33 | $C_{11}H_{20}O_4$ | 215.12888 | 0.725 | bemeth#33 | minor, $D_{11}$ |
| 215.12902 | 8.40 | $C_{11}H_{20}O_4$ | 215.12888 | 0.725 | bemeth#34 | minor, $D_{11}$ |
| 215.12900 | 8.51 | $C_{11}H_{20}O_4$ | 215.12888 | 0.545 | bemeth#35 | minor, $D_{11}$ |
| 286.16622 | 8.57 | $C_{14}H_{25}NO_5$ | 286.16600 | 0.785 | bemeth#73 | $D_{11}$ |
| 213.11345 | 8.58 | $C_{11}H_{18}O_4$ | 213.11323 | 1.014 | bemeth#322 | enriched *endo*, H/D exchange $D_5/D_6/D_7$ |
| 215.12904 | 8.61 | $C_{11}H_{20}O_4$ | 215.12888 | 0.731 | bemeth#3 | major, $D_{11}$ presumed 2*R*,3*S* |
| 229.10833 | 8.65 | $C_{11}H_{18}O_5$ | 229.10815 | 0.778 | bemeth#4 | enriched *hacl*-1, $D_9$ |
| 286.16620 | 8.69 | $C_{14}H_{25}NO_5$ | 286.16600 | 0.711 | bemeth#74 | $D_{11}$ |
| 286.16618 | 8.76 | $C_{14}H_{25}NO_5$ | 286.16600 | 0.641 | bemeth#75 | $D_{11}$ |
| 215.12901 | 8.78 | $C_{11}H_{20}O_4$ | 215.12888 | 0.616 | bemeth#32 | secondary, $D_{11}$ |
| 546.21123 | 8.80 | $C_{24}H_{38}NO_{11}P$ | 546.21097 | 0.478 | TBD | detected only in N2, $D_{12}$ |
| 231.12395 | 8.93 | $C_{11}H_{20}O_5$ | 213.12380 | 0.684 | bemeth#402 | detected only in *hacl-1(tm6725)* |
| 215.12894 | 9.01 | $C_{11}H_{20}O_4$ | 215.12888 | 0.266 | bemeth#37 | minor, $D_{11}$ |
| 361.18722 | 9.08 | $C_{17}H_{30}O_8$ | 361.18679 | 1.194 | bemeth#8 | putative glucoside |
| 591.17185 | 9.13 | $C_{22}H_{33}N_4O_{13}P$ | 591.17090 | 1.615 | gluric#421 | MOGL, enriched *endo*, co-eluting cyclopropyl isomer, $D_{11}$ |
| 215.12885 | 9.14 | $C_{11}H_{20}O_4$ | 215.12888 | 0.152 | bemeth#38 | minor, $D_{11}$ |
| 576.22211 | 9.21 | $C_{25}H_{39}NO_{12}P$ | 576.22154 | 1.003 | oglu#421 | MOGL, *endo* |
| 591.17178 | 9.26 | $C_{22}H_{33}N_4O_{13}P$ | 591.17090 | 1.506 | gluric#422 | MOGL, enriched *endo*, $D_{11}$ |
| 607.16641 | 9.28 | $C_{22}H_{33}N_4O_{14}P$ | 607.16581 | 0.984 | gluric#431 | MOGL, enriched *endo* |
| 268.15553 | 9.28 | $C_{14}H_{23}NO_4$ | 268.15543 | 0.369 | bemeth#622 | |
| 441.15345 | 9.29 | $C_{17}H_{31}O_{11}P$ | 441.15312 | 0.747 | bemeth#82 | putative phosphorylated glulcoside, *endo* |
| 558.21201 | 9.64 | $C_{25}H_{38}NO_{11}P$ | 558.21097 | 1.854 | TBD | $D_{12}$ |
| 213.11343 | 10.06 | $C_{11}H_{18}O_4$ | 213.11323 | 0.529 | bemeth#321 | H/D exchange $D_8/D_9$ |
| 199.13413 | 10.26 | $C_{11}H_{20}O_3$ | 199.13397 | 0.811 | bemeth#25 | co-eluting isomer, clearly differential in *endo*, $D_{11}$ |
| 215.12900 | 10.46 | $C_{11}H_{20}O_4$ | 215.12888 | 0.559 | bemeth#39 | minor |
| 201.14983 | 10.71 | $C_{11}H_{22}O_3$ | 201.14962 | 1.050 | bemeth#203 | minor |
| 591.17166 | 10.96 | $C_{22}H_{33}N_4O_{13}P$ | 591.17090 | 1.297 | gluric#423 | MOGL, enriched *endo*. $D_{12}$ |
| 607.16623 | 10.98 | $C_{22}H_{33}N_4O_{14}P$ | 607.16581 | 0.690 | gluric#432 | MOGL, enriched *endo* |
| 256.15555 | 11.36 | $C_{13}H_{23}NO_4$ | 256.15543 | 0.472 | bemeth#53 | minor |
| 286.16616 | 11.37 | $C_{14}H_{25}NO_5$ | 286.16600 | 0.574 | bemeth#7 | major, $D_{12}$ |
| 256.15555 | 11.42 | $C_{13}H_{23}NO_4$ | 256.15543 | 0.472 | bemeth#54 | minor, $D_{12}$ |

| | | | | | | |
|---|---|---|---|---|---|---|
| 286.16624 | 11.57 | $C_{14}H_{25}NO_5$ | 286.16600 | 0.873 | bemeth#72 | secondary, $D_{12}$ |
| 256.15535 | 11.76 | $C_{13}H_{23}NO_4$ | 256.15543 | 0.301 | bemeth#5 | major, $D_{12}$ |
| 256.15558 | 11.93 | $C_{13}H_{23}NO_4$ | 256.15543 | 0.571 | bemeth#52 | minor |
| 284.15051 | 11.95 | $C_{14}H_{23}NO_5$ | 284.15035 | 0.568 | bemeth#721 | minor |
| 268.15553 | 12.19 | $C_{14}H_{23}NO_4$ | 268.15543 | 0.365 | bemeth#621 | minor |
| 270.17116 | 12.46 | $C_{14}H_{25}NO_4$ | 270.17108 | 0.275 | bemeth#6 | major, $D_{12}$ |
| 199.13412 | 12.59 | $C_{11}H_{20}O_3$ | 199.13397 | 0.794 | bemeth#23 | unknown structure, $D_{12}$, |
| 201.14976 | 12.62 | $C_{11}H_{22}O_3$ | 201.14962 | 0.718 | bemeth#202 | minor |
| 270.17118 | 12.65 | $C_{14}H_{25}NO_4$ | 270.17108 | 0.350 | bemeth#62 | secondary, $D_{12}$ |
| 199.13408 | 12.67 | $C_{11}H_{20}O_3$ | 199.13397 | 0.580 | bemeth#22 | *2S,3S* (*anti*), minor, $D_{12}$ |
| 199.13408 | 12.83 | $C_{11}H_{20}O_3$ | 199.13397 | 0.580 | bemeth#2 | *2R,3S* (*syn*), major, $D_{12}$ |
| 197.11838 | 12.89 | $C_{11}H_{18}O_3$ | 197.11832 | 0.300 | bemeth#221 | |
| 540.20039 | 13.18 | $C_{25}H_{36}NO_{10}P$ | 540.20041 | 0.033 | iglu#201 | MOGL, enrichced *exo*, $D_{12}$ |
| 252.16065 | 13.20 | $C_{14}H_{23}NO_3$ | 252.16052 | 0.523 | bemeth#521 | |
| 201.14977 | 13.52 | $C_{11}H_{22}O_3$ | 201.14962 | 0.762 | bemeth#201 | |
| 199.13413 | 14.37 | $C_{11}H_{20}O_3$ | 199.13397 | 0.807 | bemeth#24 | unknown structure |
| 183.13928 | 26.65 | $C_{11}H_{20}O_2$ | 183.13905 | 1.239 | bemeth#1 | Long HPLC method, post-column ion pairing |

| **ES(+) Obs. *m/z*** | **RT (min)** | **Molecular Formula** | **ES(+) Theor. *m/z*** | ***m/z* error (ppm)** | **SMID-DB #** | **Comments** |
|---|---|---|---|---|---|---|
| 260.18523 | 8.90 | $C_{13}H_{25}NO_4$ | 260.18564 | 1.576 | bemeth#101 | |
| 244.19040 | 11.56 | $C_{13}H_{25}NO_3$ | 244.19072 | 1.330 | bemeth#102 | |
| 246.20612 | 12.12 | $C_{13}H_{27}NO_3$ | 246.20637 | 1.022 | bemeth#103 | enriched in *hacl-1(tm6725)* |
| 246.20606 | 12.36 | $C_{13}H_{27}NO_3$ | 246.20637 | 1.272 | bemeth#104 | detected only in *hacl-1(tm6725)* |
| 230.21103 | 12.98 | $C_{13}H_{27}NO_2$ | 230.21146 | 1.689 | bemeth#105 | |

**Supplementary Table 3.** Primers used for genotyping.

| Strain | Primer Sequence |
|---|---|
| FCS7 *hacl-1(tm6725)* II | Fwd: GAAGTAGGAATGGCAGCACAAG |
| | Rev: GGCACTGCTGAACTTGTGTAGCTC |
| | Int. Fwd: CTGCTGGCCTGTAGTCTGTATTG |
| FCS40 *fcmt-1(gk155709)* II | Fwd: CCGAGCACTCTGGGAGATTG |
| | Rev: GTGCTCACCAAATCCCACCG |
| FCS20 *fcmt-1(tm2382)* II | Fwd: CATCCAGGCGCTGGAATTC |
| | Rev: GCTCAATCGAAACCCGTGC |

**Supplementary Table 4.** Primers used for gene expression analysis by RT-PCR.

| Gene | Primer Sequence |
|---|---|
| *act-1* | Fwd: ACGACGAGTCCGGCCCATCC |
| | Rev: GAAAGCTGGTGGTGACGATGGTT |
| *fat-7* | Fwd: GGAAGGAGACAGCATTCATTGCG |
| | Rev: GTCTTGTGGGAATGTGTGGTGG |
| *fat-6* | Fwd: GGAAATTGTGTGGCGTAACG |
| | Rev: GTATGATTTGTGGGACCAGAGACG |

**Supplementary Table 5.** $^1$H NMR spectroscopic data of natural becyp#2, methanol-$d_4$ (800 MHz, $CD_3OD$).

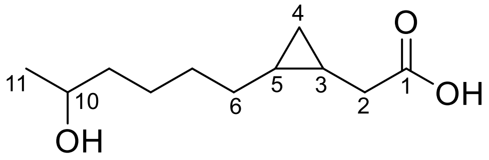

becyp#2

| **Position** | **Proton** | **$^1$H chemical shift [ppm]** | **[$^1$H,$^1$H]-Coupling constants [Hz]** |
|---|---|---|---|
| 1 | | | |
| 2 | 2-$H_a$<br>2-$H_b$ | 2.16<br>2.28 | $J_{2\text{-Ha},2\text{-Hb}}$ = 15.8, $J_{2\text{-Ha},2\text{-}3}$ = 7.8,<br>$J_{2\text{-Hb},3}$ = 6.0, |
| 3 | 3-H | 1.09 | |
| 4 | 4-$H_a$<br>4-$H_b$ | -0.15<br>0.68 | |
| 5 | 5-H | 0.78 | |
| 6-9 | 6-9H | 1.3-1.5 | |
| 10 | 10-H | 3.72 | $J_{9,10} \approx J_{10,11}$ = 6.1 |
| 11 | 11-H | 1.14 | $J_{10,11}$ = 6.1 |

**NMR spectra of synthetic compounds**

*cis*-3,4-methylenedecanoic acid, becyp#1, $^1H$ NMR spectrum (500 MHz, $CDCl_3$)

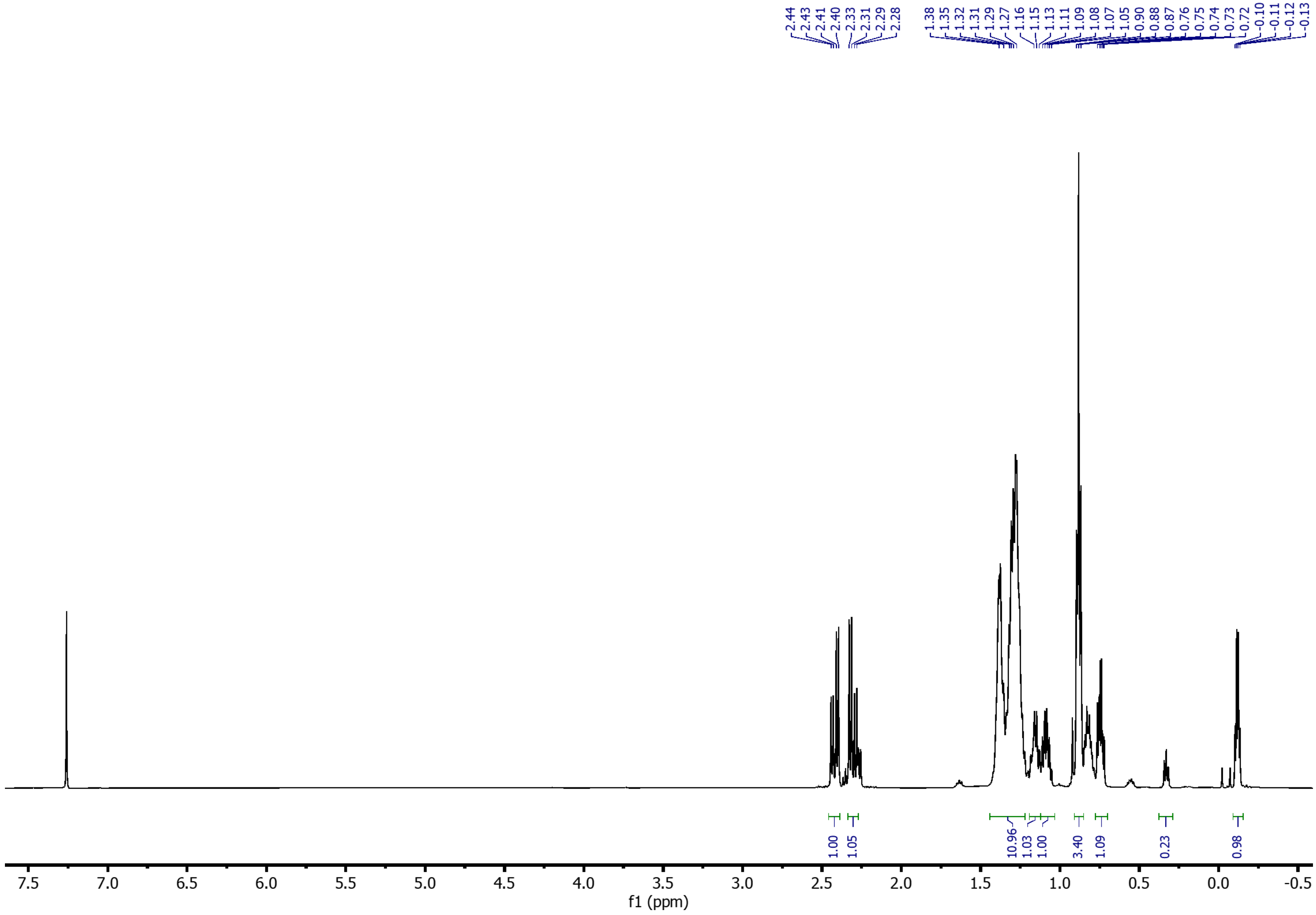

*cis*-3,4-methylenedecanoic acid, becyp#1, $^{13}C$ NMR spectrum (125 MHz, $CDCl_3$)

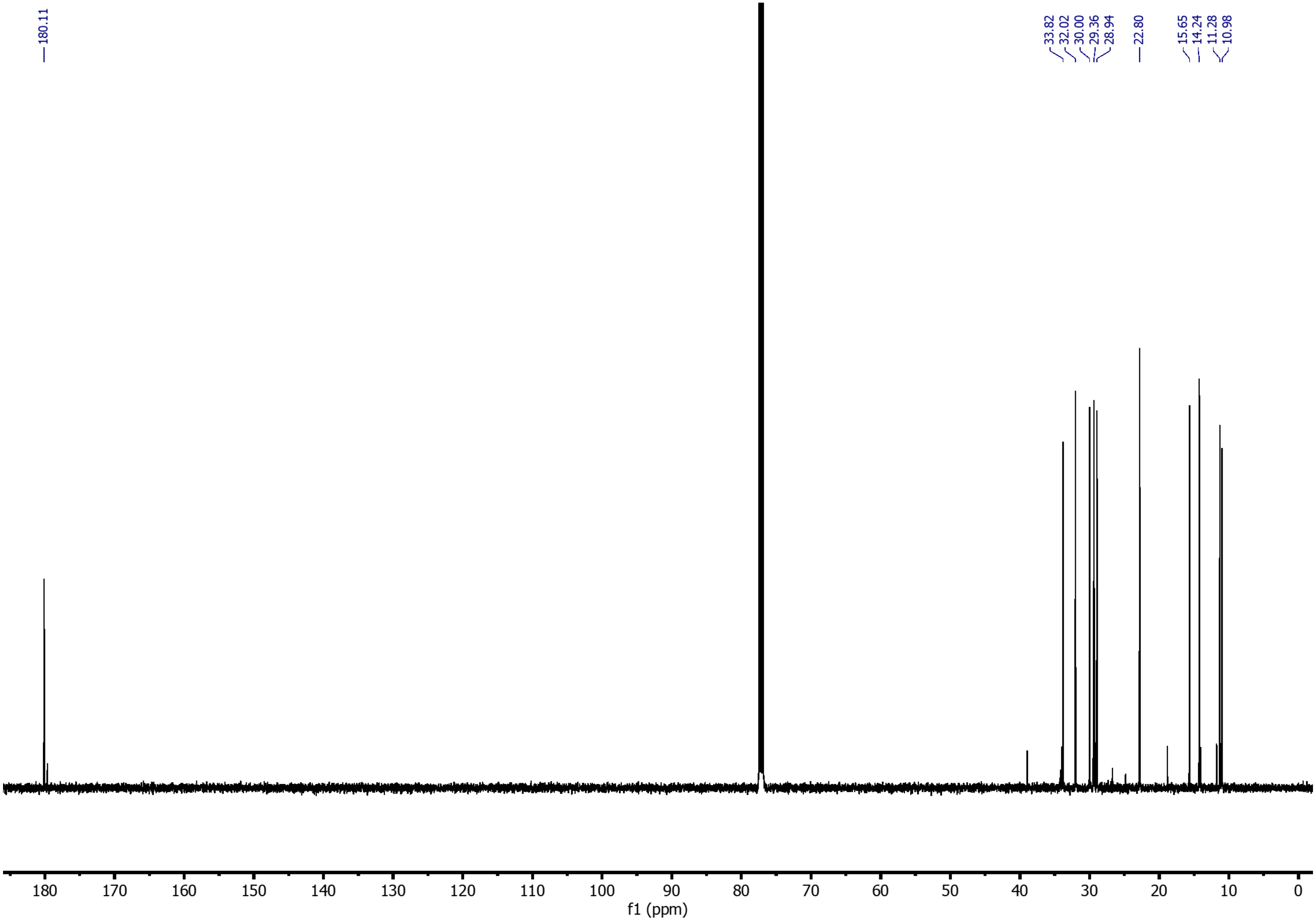

*(E)*-4-Hydroxy-non-2-ene (**5**), $^{1}$H NMR spectrum (500 MHz, $CDCl_3$)

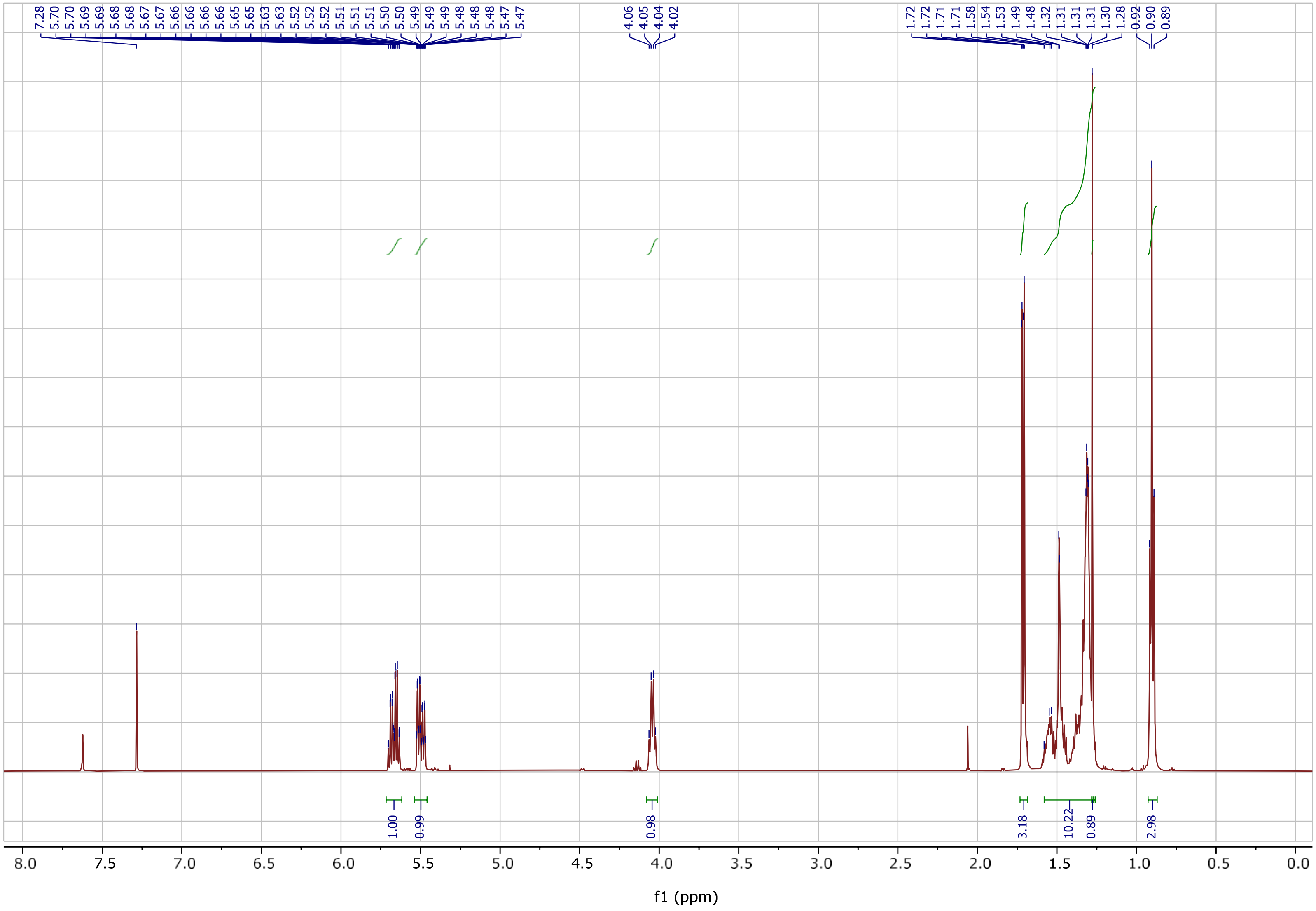

*(E)*-4-Hydroxy-non-2-ene (**5**), $^{13}$C NMR spectrum (125 MHz, $CDCl_3$)

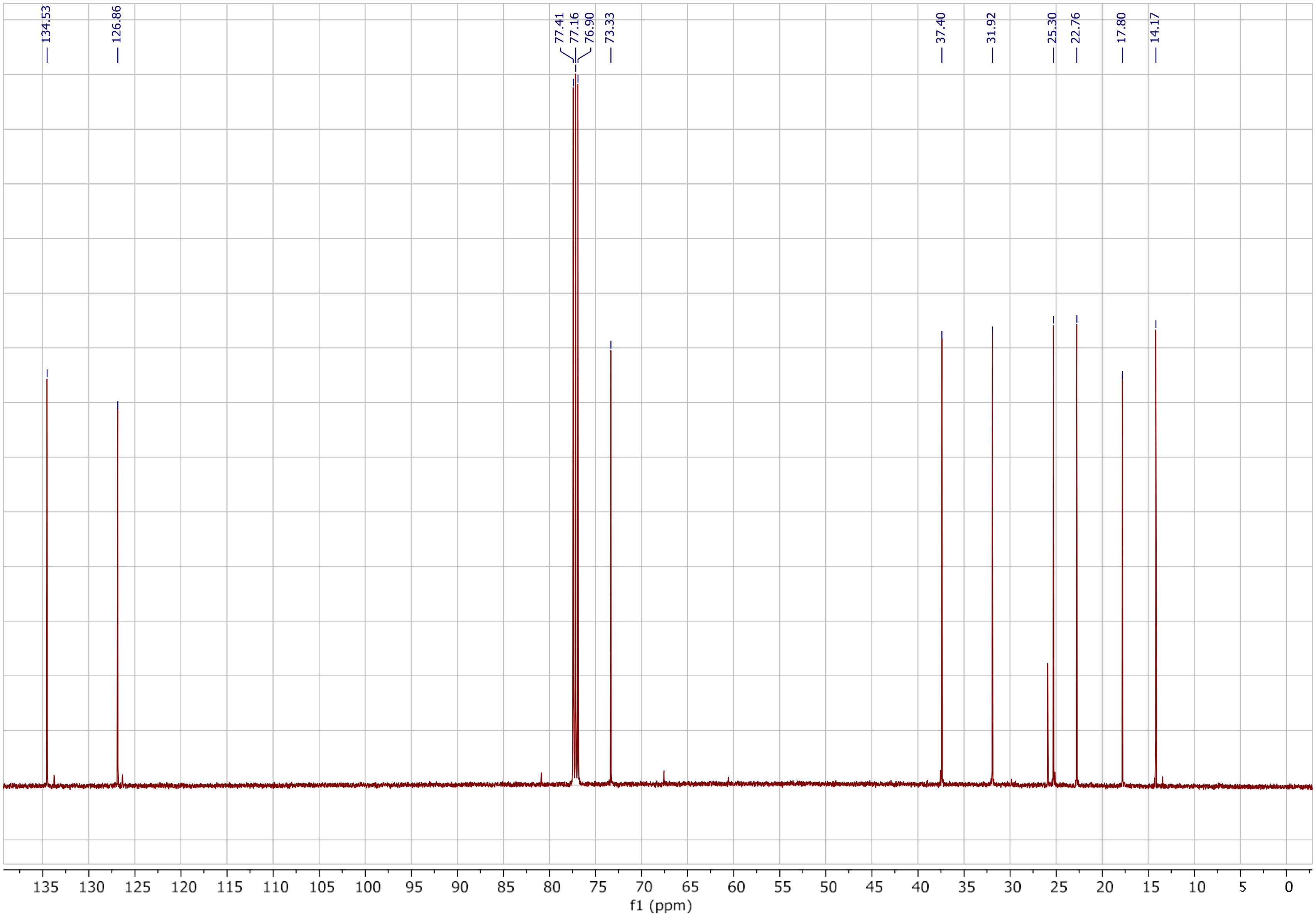

(*S,E*)-4-(Vinyloxy)-non-2-ene (**6**), $^{1}$H NMR spectrum(500 MHz, $CDCl_3$)

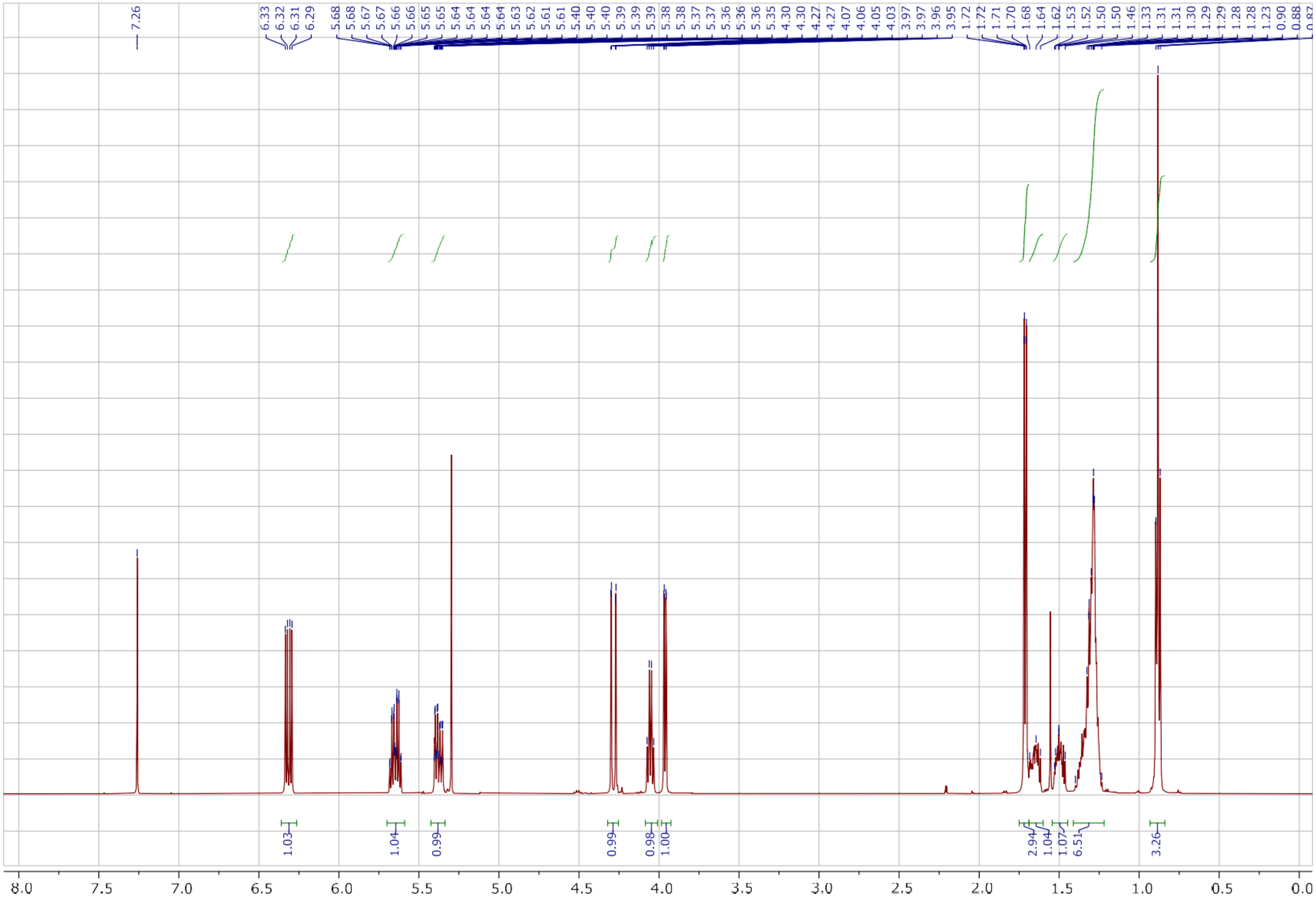

*(S,E)*-4-(Vinyloxy)-non-2-ene (**6**), $^{13}C$ NMR spectrum (125 MHz, $CDCl_3$)

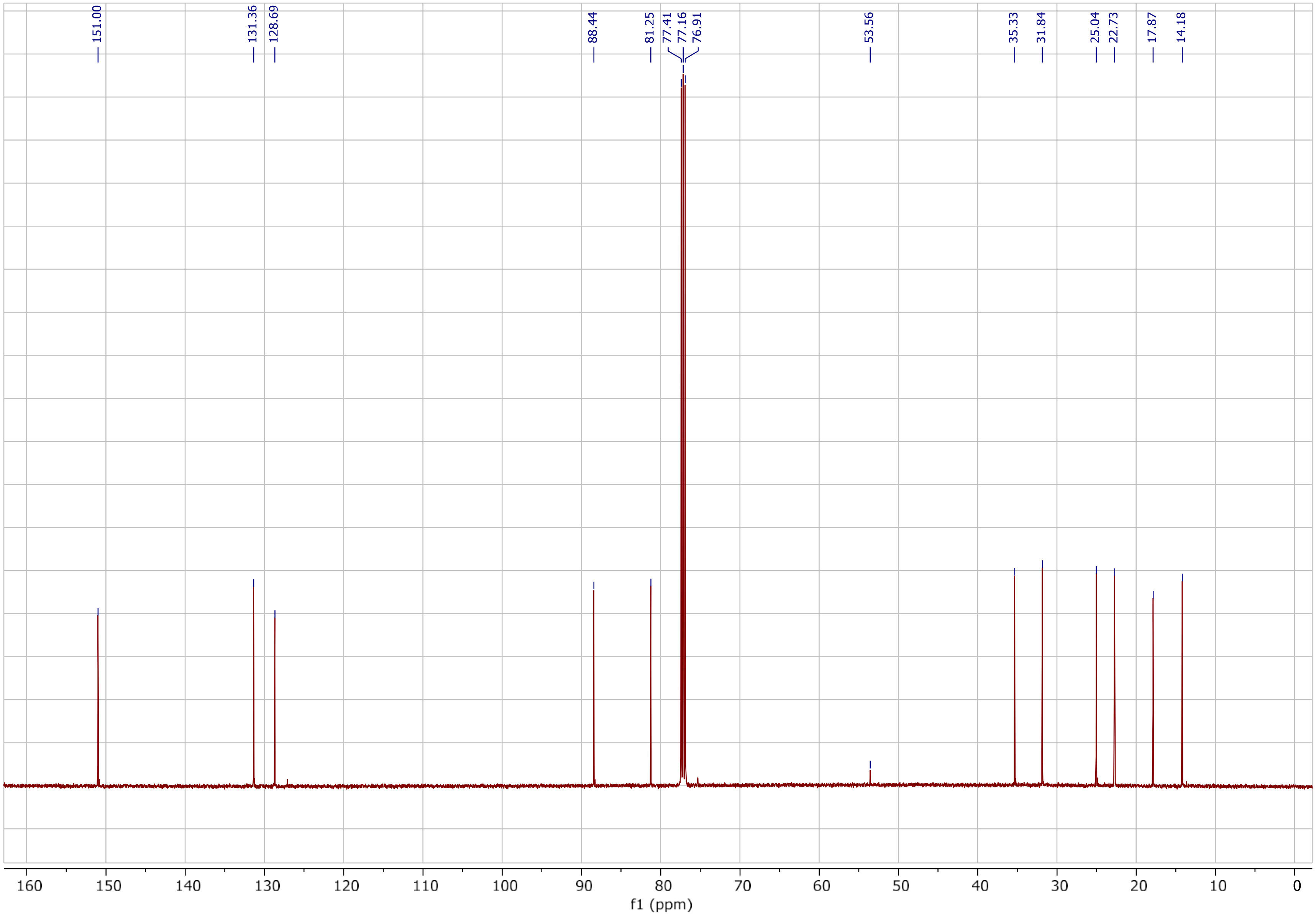

(*R,E*)-3-Methyl-dec-4-enal (**7**), $^{1}$H NMR spectrum (500 MHz, $CDCl_3$)

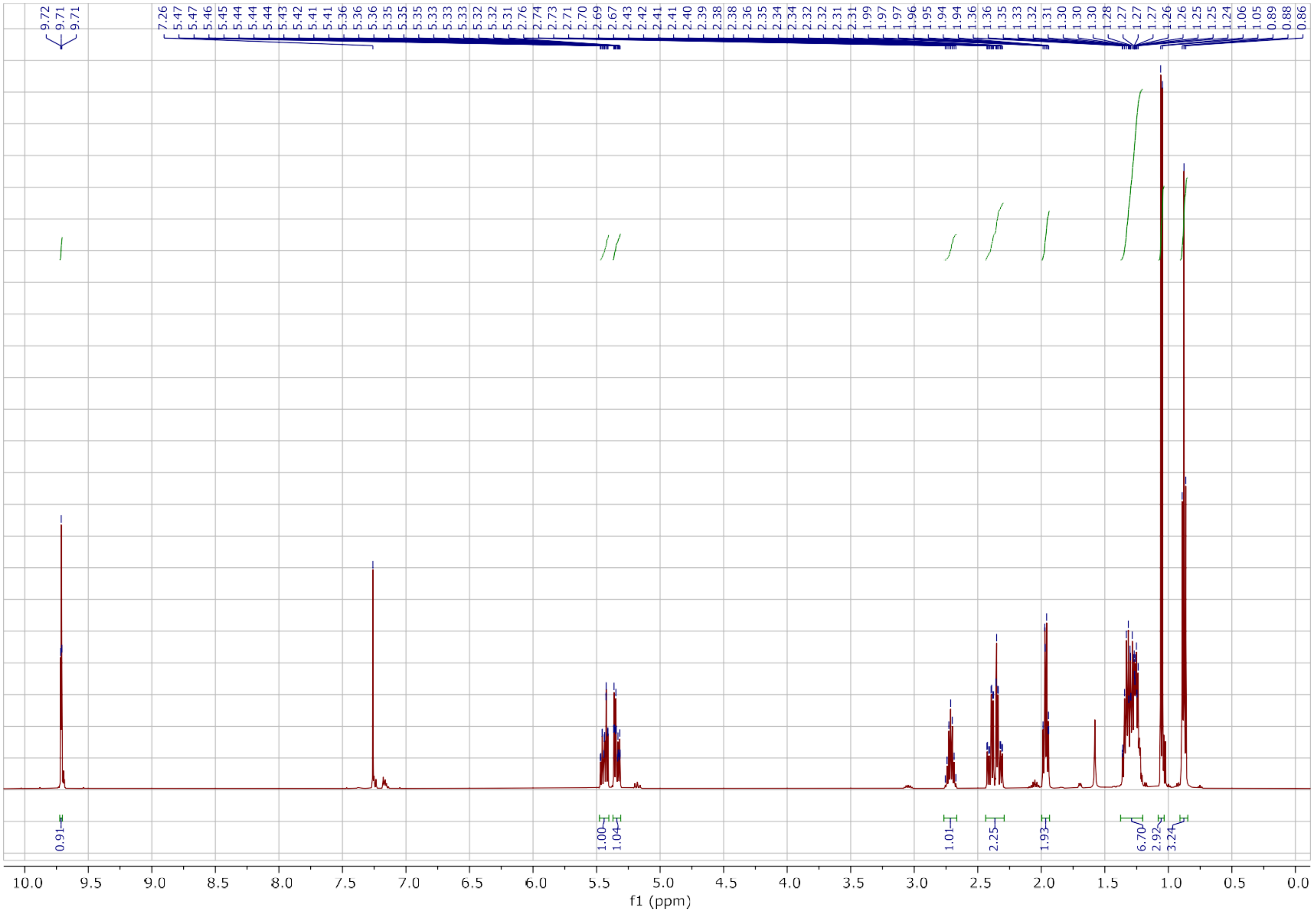

(*R,E*)-3-Methyl-dec-4-enal (**7**), $^{13}$C NMR spectrum (125 MHz, $CDCl_3$)

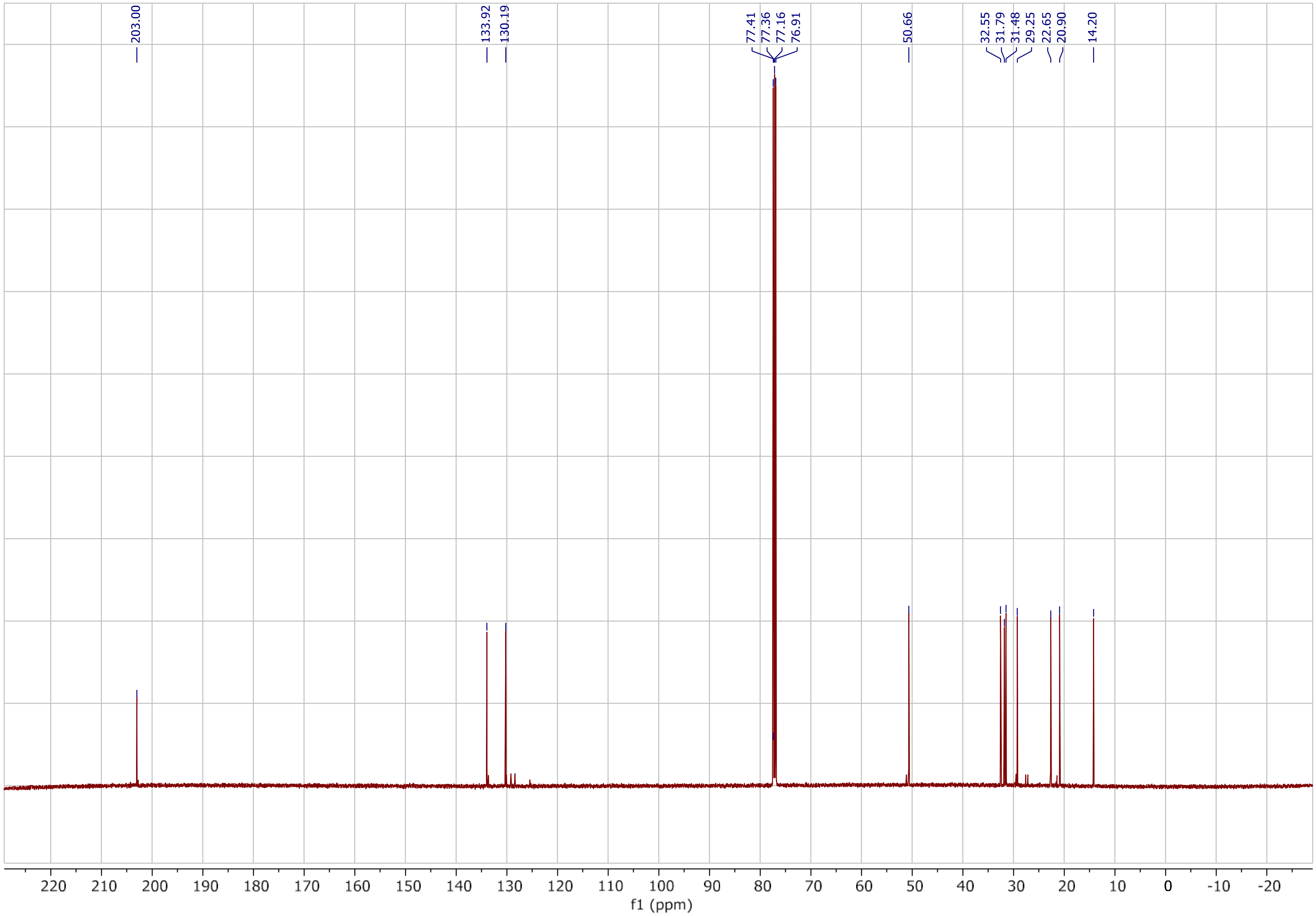

((*R,E*)-3-Methyl-dec-4-enoic acid, (*R*)-bemeth#1, $^{1}$H NMR spectrum (500 MHz, $CDCl_3$)

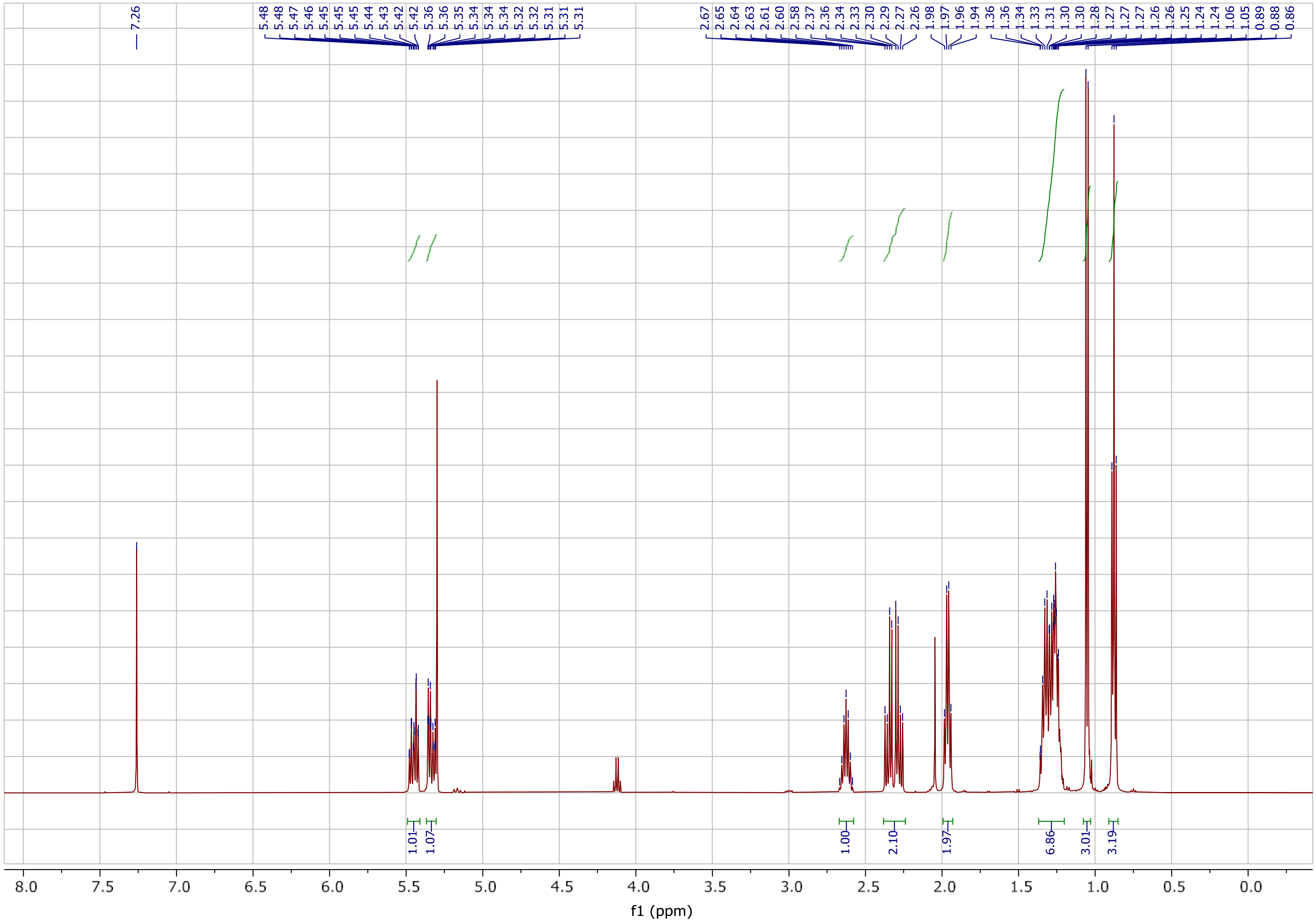

(*R,E)*-3-Methyl-dec-4-enoic acid, (*R*)-bemeth#1, $^{13}$C NMR spectrum(125 MHz, $CDCl_3$)

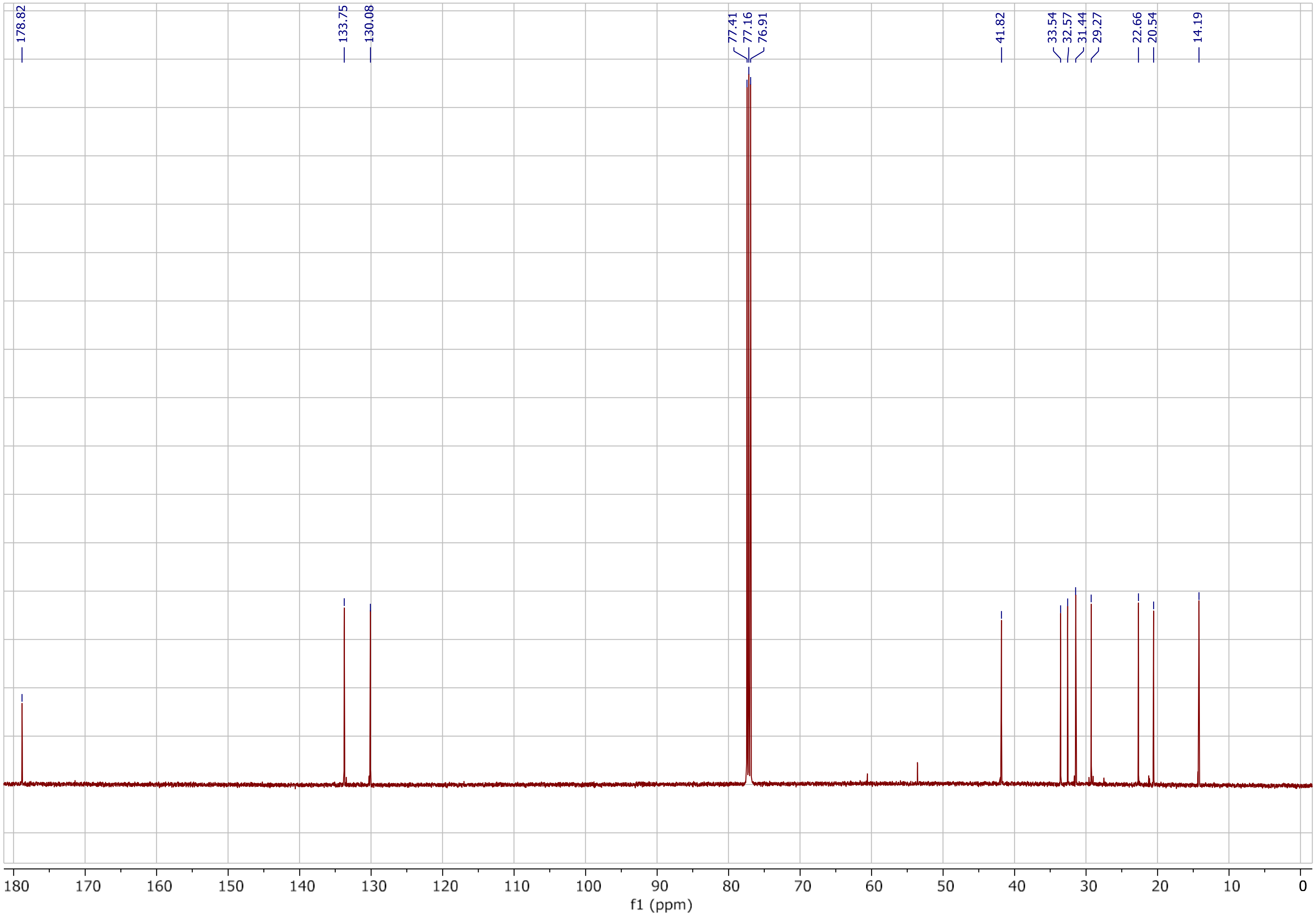

Methyl-*(R,E)*-3-methyl-dec-4-enoate (**8**), $^1$H NMR spectrum (500 MHz, $CDCl_3$)

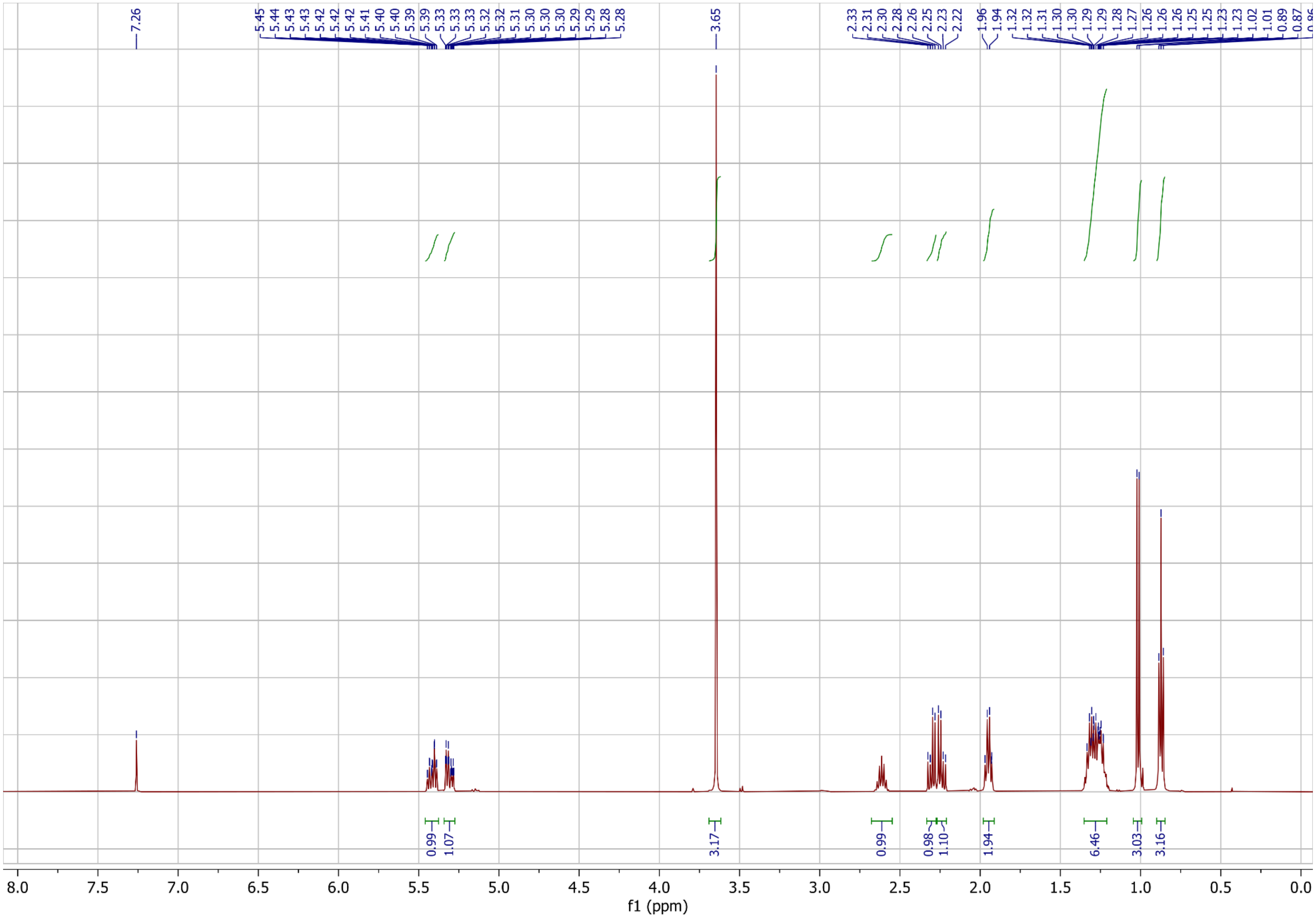

Methyl-*(R,E)*-3-methyl-dec-4-enoate (**8**), $^{13}$C NMR spectrum (125 MHz, $CDCl_3$)

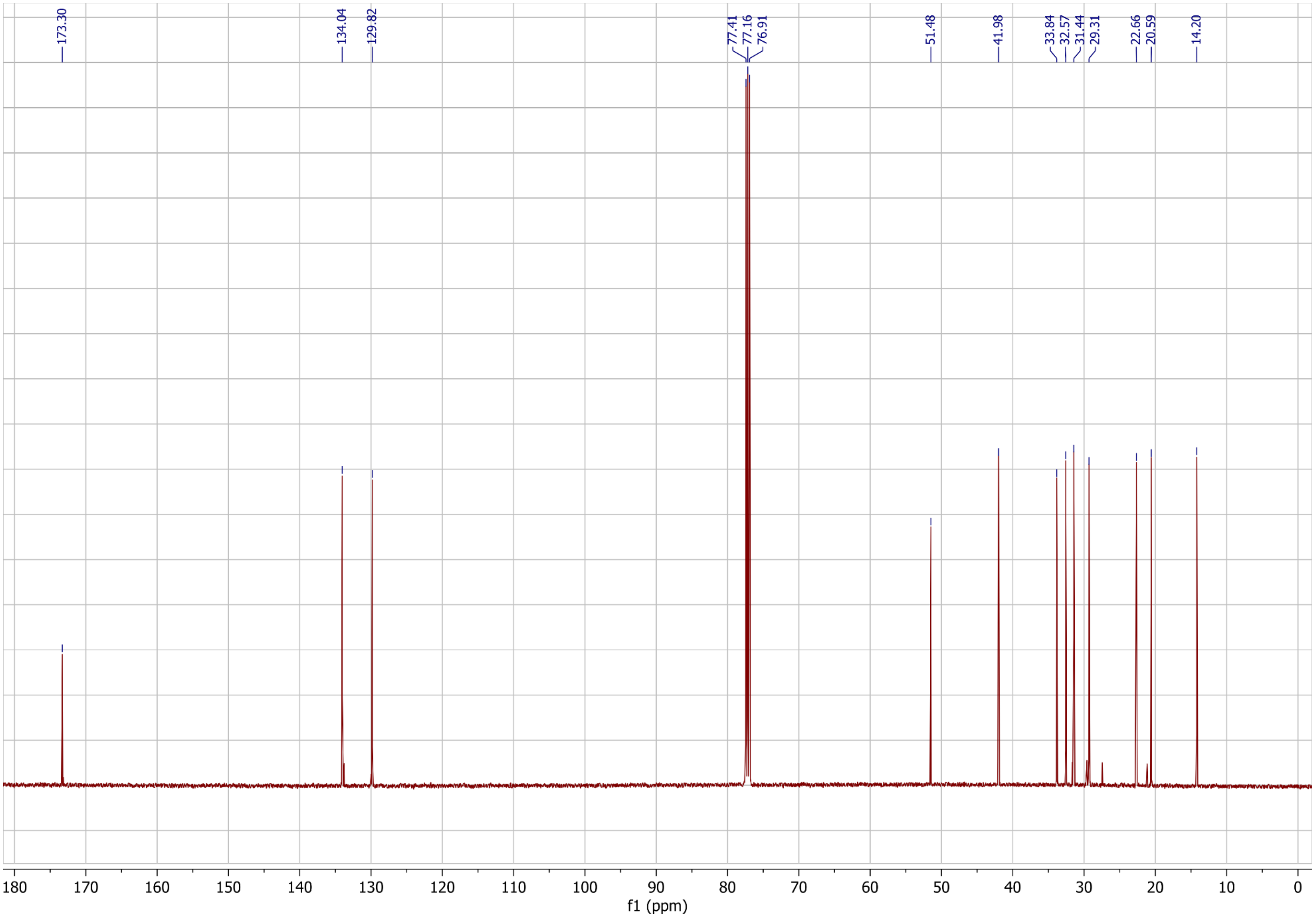

Methyl-*(2R,3S,E)*-2-hydroxy-3-methyl-dec-4-enoate (**9**), $^{1}$H NMR spectrum (500 MHz, $CDCl_3$)

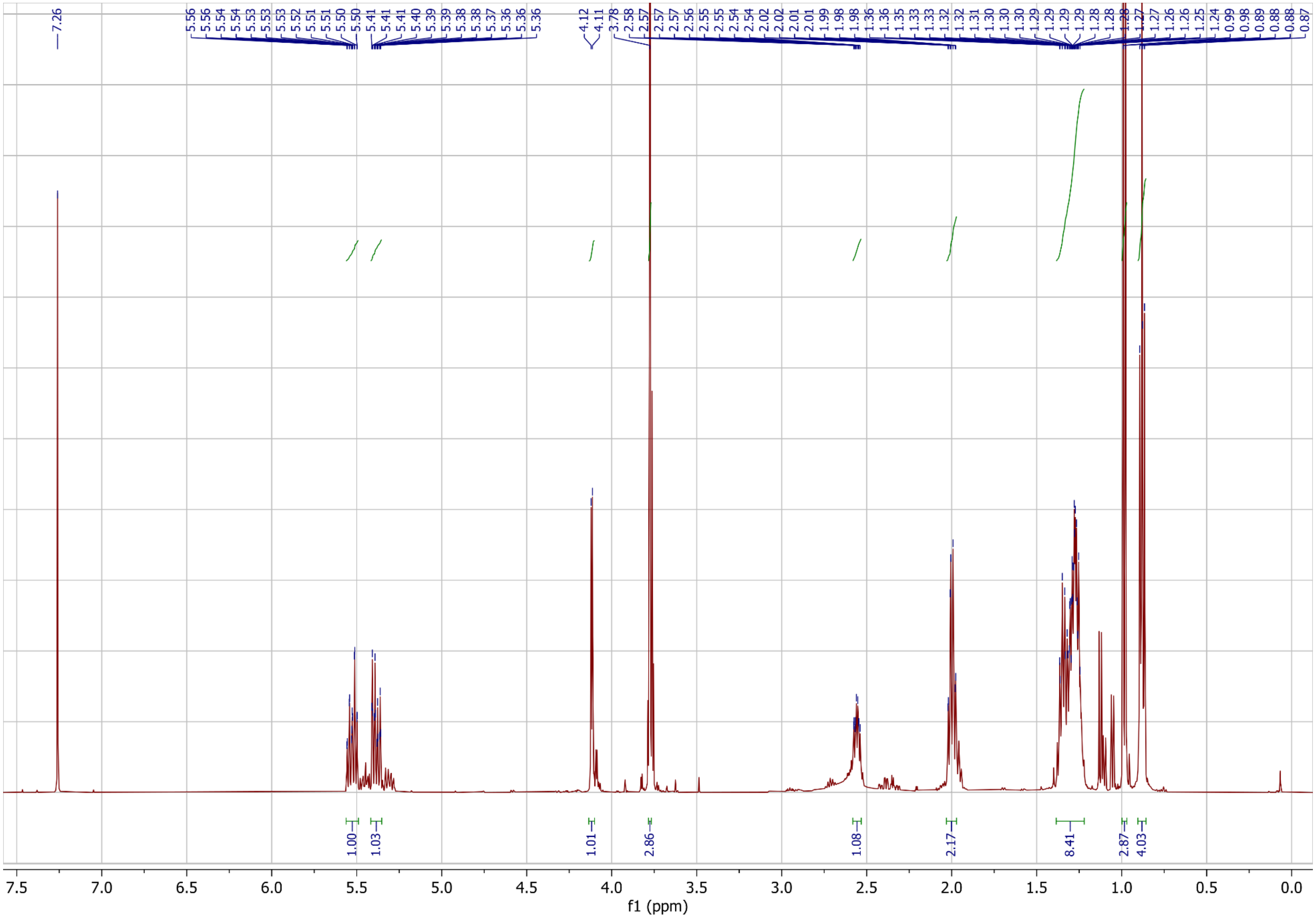

Methyl-*(2R,3S,E)*-2-hydroxy-3-methyl-dec-4-enoate (**9**) , $^{13}$C NMR spectrum (125 MHz, $CDCl_3$)

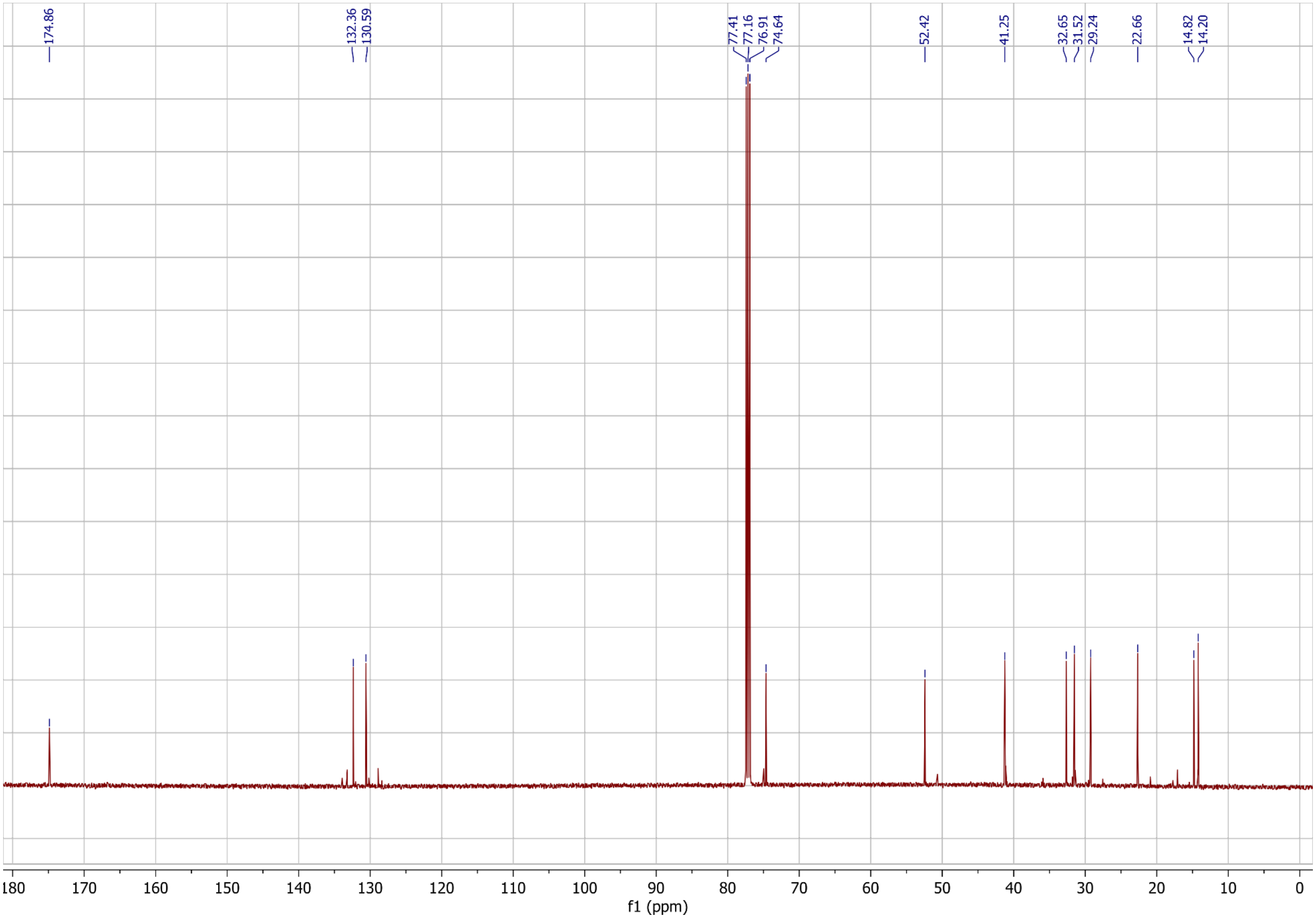

(*2R,3S,E)*-2-hydroxy-3-methyl-dec-4-enoic acid, (*2R,3S*)-bemeth#2, $^{1}$H NMR spectrum (500 MHz, $CDCl_3$)

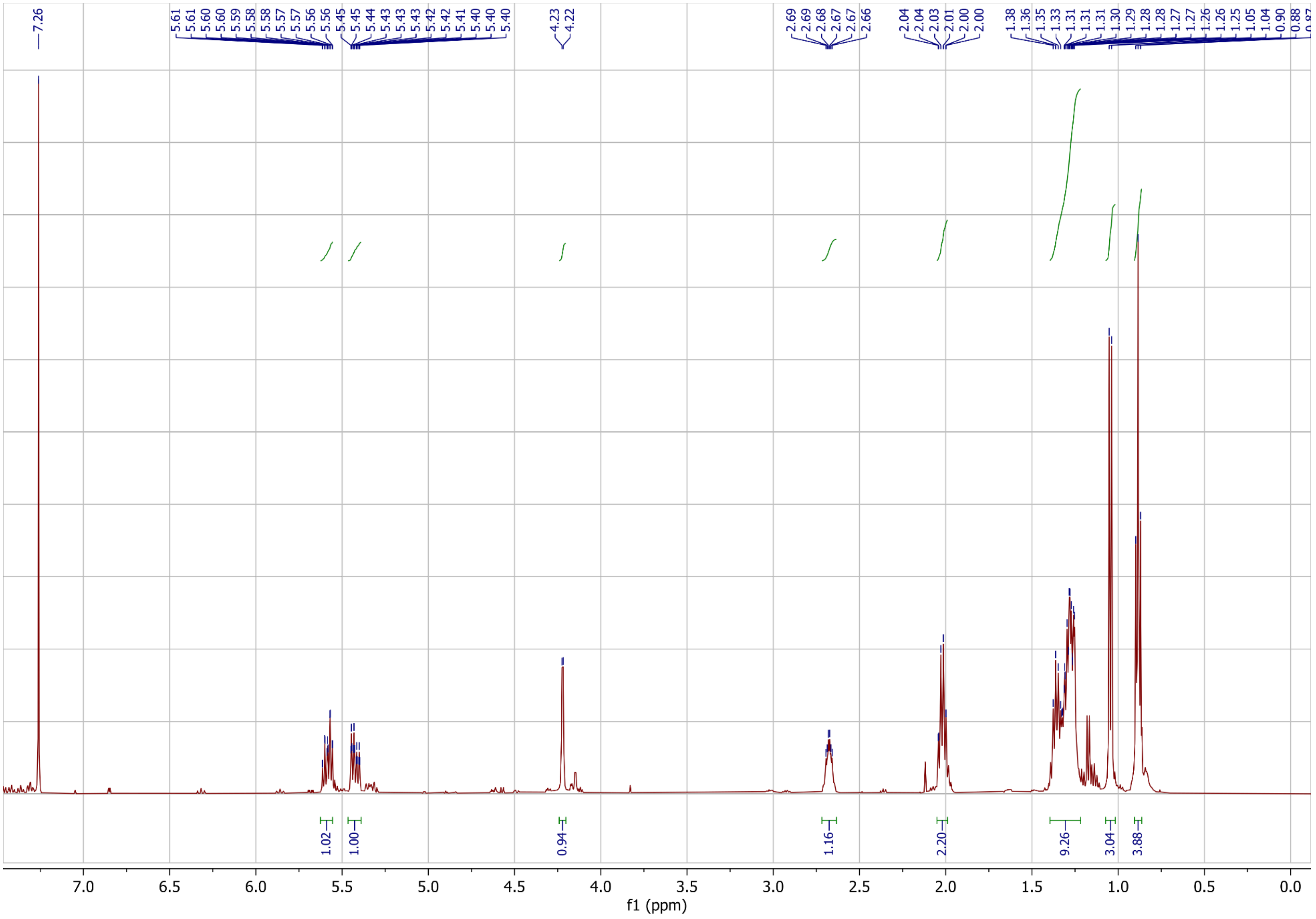

(*2R,3S,E)*-2-hydroxy-3-methyl-dec-4-enoic acid, (*2R,3S*)-bemeth#2, $^{13}C$ NMR spectrum (125 MHz, $CDCl_3$)

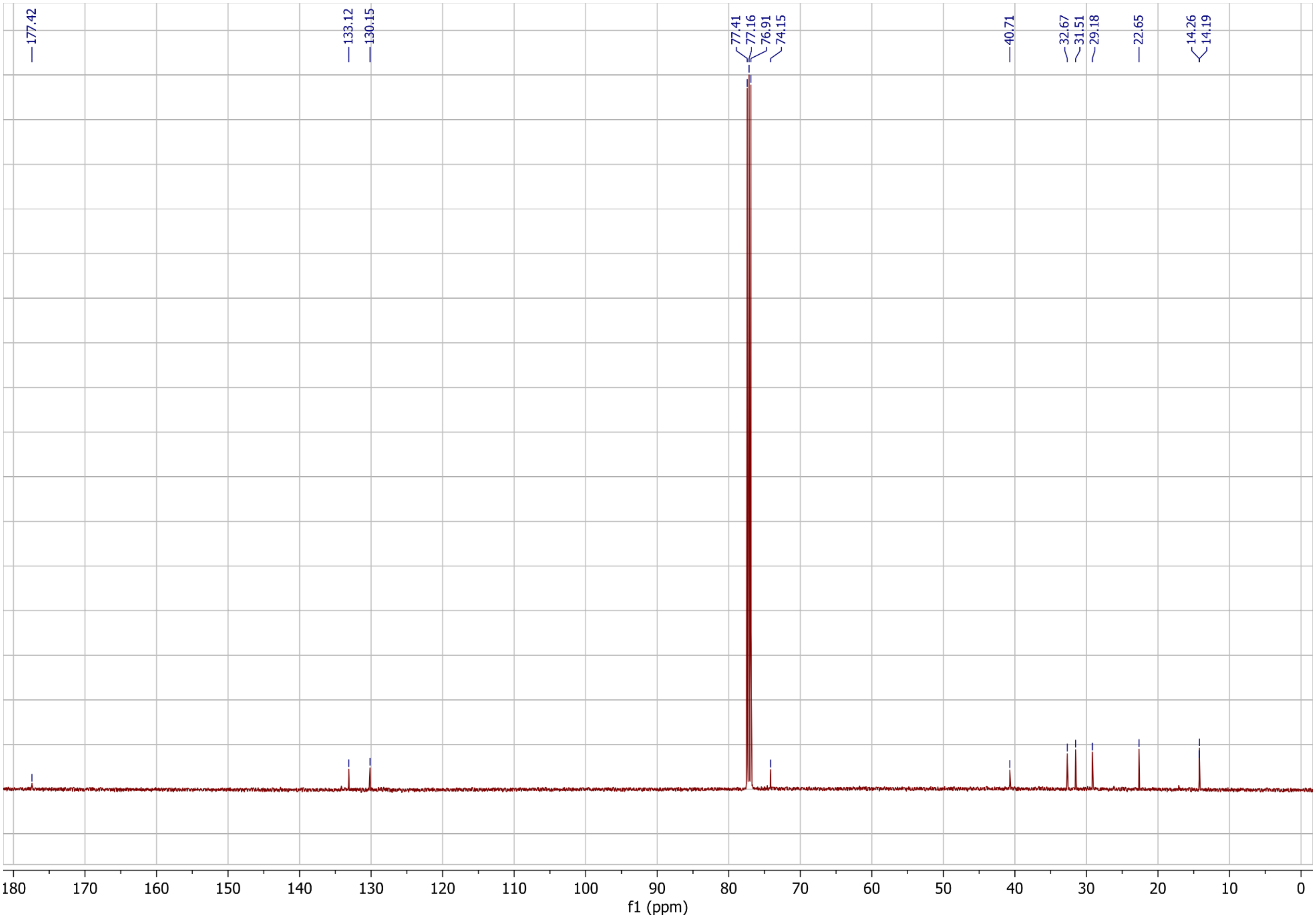

Comparison of [1]H NMR (400 MHz, $CD_3OD$) spectra for Mosher analysis. Earlier and later eluting diastereomers were separated following derivatization with (*R*)-MTPA-Cl. Derivatization of the major natural isomer, (*2R,3S*)-bemeth#2, exhibited identical chromatographic retention and chemical shifts as the later eluting fraction.

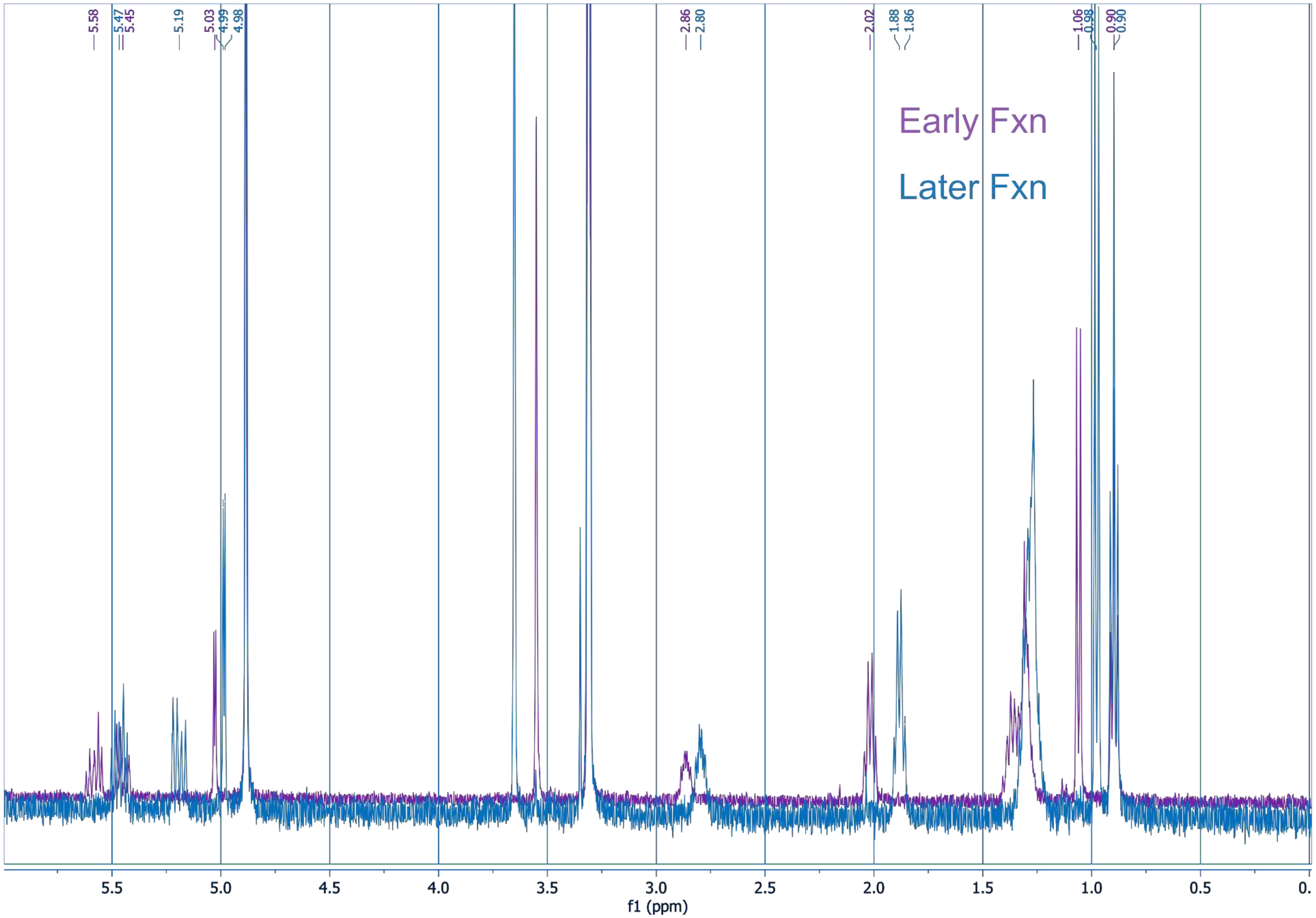
